# Counter-gradient variation in gene expression between fish populations facilitates colonization of low-dissolved oxygen environments

**DOI:** 10.1101/2023.11.14.567098

**Authors:** Janay A. Fox, David A. G. A. Hunt, Andrew P. Hendry, Lauren J. Chapman, Rowan D. H. Barrett

## Abstract

The role of phenotypic plasticity during colonization remains unclear due to the shifting importance of plasticity across timescales. Over time, genetic responses can reduce plasticity such that species in a novel environment show higher levels of plasticity than those with a longer evolutionary timescale in the environment. Therefore, comparing species in the early stages of colonization to long-established species provides a powerful approach for uncovering the role of phenotypic plasticity during different stages of colonization. We compared gene expression between the cyprinid fish *Enteromius apleurogramma*, a species that has undergone a recent range expansion, and *E. neumayeri*, a long-established native species in the same region, caught from low-dissolved oxygen (DO) and high-DO habitats. We sampled tissue either immediately after capture from the field or after a two-week acclimation under high-DO conditions, allowing us to test for both evolved and plastic differences in low-DO vs high-DO populations of each species. We found that most genes showing evolved differences in gene expression did not overlap with those showing plastic differences in gene expression. However, in the genes that did overlap, there was counter-gradient variation such that plastic and evolved gene expression responses were in opposite directions in both species. Additionally, *E. apleurogramma* had higher levels of plasticity and evolved divergence in gene expression between field populations. We suggest that the higher level of plasticity and counter-gradient variation may have allowed rapid genetic adaptation in *E. apleurogramma* and facilitated colonization. This study shows how counter-gradient variation may impact colonization of divergent oxygen environments.

## Introduction

Populations are increasingly faced with drastic shifts in their environment due to human activity and climate change (Chen et al., 2011; O’Hara et al., 2021; Yan et al., 2021). These environmental shifts may result in existing phenotypes not being well suited for current conditions, meaning that organisms must either move to more suitable habitat and/or shift their phenotypes to avoid extirpation (Parmesan & Yohe, 2003). Phenotypes can shift to new optima through adaptive genetic change, termed “evolutionary rescue” (Bell, 2013, 2017; Carlson et al., 2014); however, populations may be unable to persist long enough for evolutionary rescue to occur (Bell, 2013). Phenotypic plasticity, broadly defined as the ability of a single genotype to produce different phenotypes depending on the environment (West-Eberhard, 2003), allows for rapid phenotypic change in response to environmental conditions. Plasticity has been suggested to play a major role in the colonization of new environments (Bilandžija et al., 2020; Walter et al., 2022; Wang & Althoff, 2019; Yeh & Price, 2004), range expansions (Doudová-Kochánková et al., 2012; Otaki et al., 2010; Zarco-Perello et al., 2022), responses to climate change (Charmantier et al., 2008; Franks et al., 2014; Potts et al., 2021), and invasive species’ ability to invade (Jardeleza et al., 2022; Liao et al., 2016; Pichancourt & Klinken, 2012).

Understanding how phenotypic plasticity facilitates population persistence during colonization and range expansion is important for predicting species responses to environmental change. However, the role that phenotypic plasticity plays during these challenges remains unclear due to inconsistent results across studies (reviewed in Hendry, 2016). For example, two meta-analyses published in the same year that investigated levels of plasticity expressed in invasive versus non-invasive plants found conflicting results – one study indicated that invasive plants did not have higher levels of plasticity than non-invasive plants (Palacio-López & Gianoli, 2011), while the other found that invasive plants had significantly higher levels of plasticity (Davidson et al., 2011). When considering how plasticity will help species persist under climate change, some studies have found that higher plasticity led to increased persistence (Henn et al., 2018; Urban et al., 2014; Vedder et al., 2013), whereas other studies have suggested that plasticity will have a limited impact (Gill et al., 2014; Gunderson & Stillman, 2015; Kellermann et al., 2020). These inconsistent results suggest that the impact of plasticity is context dependent. This could be due to different ways that adaptive versus maladaptive plasticity alter species responses to environmental shifts. If plasticity is adaptive, it may allow organisms to persist in novel or changing environments through “plastic rescue” (Chevin et al., 2010; Kovach-Orr & Fussmann, 2013; Lande, 2009; Snell-Rood et al., 2018). Adaptive plastic phenotypic shifts can then be followed by additive genetic change in the same direction, termed ‘genetic assimilation’ (Schlichting & Wund, 2014; West-Eberhard, 2003). If the plasticity is maladaptive, plastic shifts move populations further from the optimum phenotype, which could hinder survival in a new environment. Genetic change in the opposite direction of the plasticity, termed ‘genetic compensation’, is then required to push phenotypes closer to the optimum (Grether, 2005). This leads to ‘counter-gradient variation’ where individuals from different environments display higher trait similarity in the field than when acclimated in a common environment (Conover & Schultz, 1995).

Another potential reason for these inconsistencies may lie in the shifting importance of plasticity across timescales. Both genetic assimilation and genetic compensation are subsets of ‘genetic accommodation’ whereby genetic responses can reduce plasticity either by reinforcing the adaptive plastic phenotype such that it no longer needs to be environmentally induced or by reversing the maladaptive plastic phenotype so that it is no longer expressed (Grether, 2005; Waddington, 1942; West-Eberhard, 2003). Thus, one could predict that species that are new to an environment would show higher levels of plasticity than those that have been in the environment for a longer evolutionary timescale and have had time for genetic accommodation to take effect. This shifting importance of plasticity through time becomes an issue when comparing native species to invading species that have already become well-established in the novel environment, as is done in many studies due to difficulties in capturing initial range expansions or colonization events. It would be more informative to compare levels of plasticity between species that differ in their experience with an environment (i.e., newly invading vs native species).

This study takes advantage of a system in which fish communities have undergone recent range shifts in response to changing environmental conditions to compare plasticity across two different time scales – very recent range-expanding versus long-established populations. Long-term monitoring of the Mpanga River drainage in Kibale National Park, Uganda has captured the range expansion of the cyprinid *Enteromius apleurogramma* northwards into the Rwembaita Swamp System (RSS), which includes a low dissolved oxygen (DO) swamp and high-DO tributary streams. Monitoring of the RSS since 1990 indicated that the system hosted only two native fishes, the cyprinid *Enteromius neumayeri* and the air-breathing catfish *Clarias liocephalus*, both of which occur in low-DO and high-DO environments. *E. apleurogramma* was first recorded in the RSS in 2015 but has since spread throughout the entire swamp and associated streams (Hunt et al., 2023). It is one of three native fish species known to have expanded their range northward in the Mpanga River system, the others being the cyprinodontid *Platypanchax modestus* (appeared in 2012) and the cichlid *Pseudocrenilabrus multicolor* (appeared in 2022). In *E. neumayeri*, there is strong phenotypic divergence between low-DO (swamp) and high-DO (stream) sites, with swamp-dwelling populations characterized by greater tolerance to hypoxia (Chapman, 2007; Olowo & Chapman, 1996), higher hematocrit (Chapman, 2007; Martinez et al., 2004), higher liver LDH activities, and higher glycolytic capacity (Chapman, 2015; Martínez et al., 2011). Populations do exchange some migrants between high DO and low DO habitats (Chapman et al., 1999; Harniman et al., 2013); however, a combination of long-term acclimation (Martínez et al., 2011), and genetic studies (Chapman et al., 1999; Harniman et al., 2013) suggest genetic differentiation between oxygen regimes over small spatial scales. *E. neumayeri* and *E. apleurogramma* inhabit very similar habitats, are phylogenetically closely related (Ndeda, 2018), and display similar patterns of divergence across DO gradients (Hunt et al., 2023). Therefore, this study system allows us to compare levels of plasticity between two similar species that have different time scales of experience with the habitat: one experiencing a recent range shift (began two years prior to sampling) and another that has a much longer evolutionary history within the area.

This range expansion of *E. apleurogramma* was likely enabled by a recent increase in temperature in the RSS that made it more similar in temperature to the original habitat of *E. apleurogramma* (Hunt et al., 2023). It is expected that the colonizing *E. apleurogramma* individuals originated from high-DO populations based on the most direct route, however it is possible that some individuals have previous experience with low-DO environments. Hypoxia, defined as DO levels under 2-3 mg O^2^/L (Vaquer-Sunyer & Duarte, 2008), is common in the heavily vegetated papyrus swamp of the RSS due to lower flow and higher rates of decomposition. DO levels can be especially limiting in aquatic environments and impose strong selective pressures on fish species (Chapman, 2015). Accordingly, variation in DO levels shape species ranges and result in many different behavioural and physiological adaptations (Nikinmaa & Rees, 2005; Richards, 2009, 2011). Due to the strong pressure that DO levels exert, it is likely that phenotypic plasticity in traits underlying hypoxia tolerance facilitated the colonization of *E. apleurogramma* (Crispo & Chapman, 2010).

Gene expression connects genotypes to phenotypes; therefore, plasticity in gene expression can serve as a link between environmental change and adaptive phenotypic plasticity (Rivera et al., 2021; Schlichting & Wund, 2014). Gene expression plasticity has been found to allow species to cope with variable environments, including hypoxia (Gracey et al., 2001; Nikinmaa & Rees, 2005; Storz et al., 2010), and to facilitate the colonization of new environments (Bittner et al., 2021; Morris et al., 2014). Hypoxia induced plasticity in gene expression can occur very rapidly – goby fish (*Gillichthys mirabilis*) exposed to hypoxia showed shifts in gene expression within eight hours that were maintained for at least six days (Gracey et al., 2001). While all genes likely display a level of plasticity in their expression, the magnitude and direction of gene expression plasticity can be compared across populations and different environments to reveal potential differences in levels of phenotypic plasticity. Additionally, some differences in gene expression have a heritable basis that selection can act on (Crawford & Oleksiak, 2007; Whitehead & Crawford, 2006). Therefore, gene expression can be involved in both plastic and evolutionary divergence. In this experiment, we compared gene expression between *E. apleurgramma*, representing a range expanding species, and *E. neumayeri*, representing a native species with evolutionary history in the area, caught from low-DO and high-DO habitats within the RSS. We sampled tissue either immediately after capture or after a two-week acclimation period at high-DO. This allowed us to test for both evolved and short-term plastic differences in low-DO vs high-DO populations of each species. We hypothesized that plasticity in gene expression underlying hypoxia tolerance facilitated the colonization of *E. apleurogramma* into divergent oxygen environments within the RSS. We predicted that gene expression would differ between colonizing and native populations such that in low-DO vs high-DO comparisons, colonizers exhibit primarily plastic gene expression whereas native populations exhibit lower plasticity but more evolved divergence due to inherited differences in gene expression. We assumed that evolved divergence between habitats reflects adaptive change, and as such, if this plasticity is maladaptive, we predicted that gene expression plasticity and evolved divergence would occur in opposite directions. For example, a plastic difference is reflected by upregulation of a gene in low-DO relative to high-DO conditions, whereas the evolved difference is reflected by downregulation of that gene in low-DO relative to high-DO conditions. In contrast, if plasticity is adaptive, gene expression plasticity and evolved divergence should occur in parallel directions.

## Methods

### Ethics statement

Permission to carry out this work came from the Uganda National Council for Science and Technology, the Uganda Wildlife Authority, and McGill University Animal Care (AUP 5029).

### Study site

This study was conducted within the Rwembaita Swamp system (RSS) (00.58875 °N 030.37222 °E) in Kibale National Park, Uganda. In this papyrus (*Cyperus papyrus*) swamp, low water flow and mixing combined with high input of organic matter and levels of shade result in low dissolved oxygen levels, averaging 0.99 mg/L between 1993 and 2019 (for DO data see (Chapman et al., 2022)). However, the swamp has associated streams and river tributaries where increased flow and turbulence leads to much higher average dissolved oxygen levels (∼6 mg/L). Between 1994 and 2016, average local air temperatures have increased 1.45 °C and concordantly average water temperatures have increased by 1.41 °C (Lauren Chapman, unpublished data). It is possible that this shift in temperature has facilitated the expansion of *E. apleurogramma* into the swamp as historical populations reside in locations approximately 200 m lower in elevation that would have a predicted average temperature that is 1.3 °C higher than the RSS (Hunt et al., 2023), although actual temperature records for the lower site are not available.

### Fish collection and acclimation trials

Similarly sized adult *E. neumayeri* (average standard length (SL): 5.97 cm +/- 1.05 cm) and *E. apleurogramma* (average SL: 4.23 cm +/- 0.28) were collected from a swamp and stream pair separated by ∼200 m. Collections were done on June 6^th^, 2017 using barrel minnow traps with a mesh size of 6.35 mm and throat openings of 25.4 mm baited with bread. Fish were randomly divided into two categories: those to be immediately sacrificed and those to be acclimated for two weeks. Fish selected for immediate sacrifice were euthanized using clove oil within 10 minutes of being pulled from the trap; gills were then extracted as quickly as possible and placed into RNAlater (Qiagen, Hilden, Germany). Gill tissue was chosen due to the central role it plays in respiration and its known plasticity in response to different DO levels (Sollid & Nilsson, 2006). Samples were initially stored at ambient temperature (∼30°C) for 4-8 hours before being returned to the field station (Makerere University Biological Field Station, MUBFS) where they were stored at 4°C. Fish selected to be acclimated were returned to the field station in small containers of well-oxygenated water. Fish were marked with a subdermal dye mark just below the dorsal fin, with combinations of colour and side of body indicating population and species. Fish were held for 14 days at ambient temperature in two open air ponds (∼2 m diameter by 50 cm depth) equipped with air pumps to ensure full oxygenation of the water. Approximately 10 fish of each of the two species were held in each pond for a total of 20 fish per pond: five from each species from the hypoxic swamp and five from each species from the normoxic stream.

Pools were monitored daily for temperature and dissolved oxygen (Supplemental Table 1), and for fish morbidity and mortality. Fish in the pools were fed ad libitum, and water changes were performed every three days. Acclimated fish were sacrificed after 14 days in a manner identical to immediately sacrificed fish: euthanized by clove oil and the gills immediately extracted and placed in RNAlater. Samples were held at 4°C for 30 days before being transported at ambient temperature over a period of 36 hours to McGill University, where they were stored at -20°C until extraction. Sex is cryptic in these species therefore there are no sex data for these samples.

### RNA extraction and sequencing

DNA and RNA were extracted using AllPrep DNA/RNA Kits (Qiagen, Hilden, Germany) following the manufacturer’s protocol. We measured RNA quality using the Agilent RNA 6000 Nano Kit (Agilent, Santa Clara, United States) and quantity using the Quant-it RiboGreen RNA Assay Kit (Invitrogen, Waltham, United States). Seventy out of 80 samples were deemed of sufficient quantity and quality for sequencing (Table 1). Samples were sent to the McGill Genome Center (Montréal, Canada) for library preparation and sequencing. Libraries were prepared using the NEB Ultra II Directional RNA Library Prep Kit (New England BioLabs, Ipswich, United States), and all samples were run on one lane of the NovaSeq6000 S4 v1.5 (Illumina, San Diego, United States).

**Table 1.** Number of *E. neumayeri* (EN) and *E. apleurogramma* (EA) fish retained for analysis for each sampling category and species.

| Sampling Category | Population |  |
| --- | --- | --- |
|  | Low DO | High DO |
| Immediate | EN:8, EA:10 | EN: 9, EA: 9 |
| Acclimation | EN: 7, EA: 11 | EN: 8, EA: 7 |

### Read quality control and de novo assembly

We used Rcorrector v1.0.4 (Song & Florea, 2015)to remove erroneous k-mers using default settings and then TrimGalore! v0.6.6 (https://www.bioinformatics.babraham.ac.uk/projects/trim_galore/) to trim low quality bases (phred < 5) and adaptor contamination. Next, we generated quality reports on our samples using FastQC (Andrews, 2019) and found that there were many overrepresented sequences that corresponded to rRNA when searched using BLAST (Altschul et al., 1990). To counteract this, we constructed a rRNA database using the SSUParc and LSUParc v138.1 files in the silva database (accessed Dec 2022; www.arb-silva.de) then mapped the reads to the database using bowtie2 v2.3.4.3 (Langmead & Salzberg, 2012) with the --nofw flag for dUTP based libraries and the –very-sensitive-local preset option. Only read pairs for which neither read mapped to the database were retained for further analysis. Again, FastQC was run and read quality was checked. One *E. neumayeri* (low-DO, immediate) sample was removed due to poor sample quality.

### Trinity de novo assembly

As there is no reference genome available for either species, we performed de novo assembly for each species separately using Trinity v2.15.0 (Grabherr et al., 2011) with –SS_lib_type set to RF for dUTP based libraries and default settings. To assess the quality of our trinity assemblies we looked at several assessment metrics. First, we used bowtie2 v2.3.4.3 (Langmead & Salzberg, 2012) to align reads from each individual to its corresponding assembly and examined the RNA-seq read representation of the assembly. Next, we computed ‘gene’ contig Nx length statistics where at least x% of the assembled transcript nucleotides are found in contigs of at least Nx length for x = 10-50 along with counts of transcripts and “genes” and median contig length using a custom perl script in the Trinity toolbox (TrinityStats.pl). Then, we used BUSCO v5.2.2 (Manni et al., 2021) with the vertebrata_odb10 BUSCO set (accessed Feb 2023) using the transcriptome setting to estimate the completeness and redundancy of the assembly. To functionally annotate the assemblies, we used TransDecoder v 5.7.0 to predict coding regions (Haas, BJ. https://github.com/TransDecoder/TransDecoder). Then, we used Trinotate v4.0.0 (Bryant et al., 2017) to compare predicted coding regions and entire transcripts to established protein databases, Swiss-Prot (Bairoch & Apweiler, 1997) and PFam (Punta et al., 2012) (both accessed May 2023). We also used Infernal v1.1.4 (Nawrocki & Eddy, 2013) to search the noncoding RNA database Rfam v14.9 (accessed May 2023). Using Trinotate, we annotated coding regions for signal peptides with Signal P v6.0 (Teufel et al., 2022), transmembrane helices with tmHMM v2 (Krogh et al., 2001), and domain content with Eggnog-mapper v2 (Cantalapiedra et al., 2021).

### Quantification and differential gene expression analysis

We used *Salmon* v1.10.1 (Patro et al., 2017) to quantify transcripts and generate gene counts using the Trinity gene transcript map generated for each species. All further analyses were performed in R v4.2.2 (R Core Team, 2022). Differential gene expression was analyzed using *edgeR* v3.40.2 (Robinson et al., 2010) by running four pairwise comparisons within each species: 1. Low DO population, immediate sampling vs, low DO population, sampling after acclimation (plastic differences and lab effects; L-I vs L-A); 2. High DO population, immediate sampling vs high DO population, sampling after acclimation (lab effects; H-I vs H-A); 3. Low DO population, immediate sampling vs high DO population, immediate sampling (plastic and evolved differences; L-I vs H-I); and 4. Low DO population, sampling after acclimation vs high DO population, sampling after acclimation (evolved differences; L-A vs H-A) (Figure 1).We filtered out lowly expressed genes using a count-per-million (CPM) threshold of one, corresponding to a count of six reads in the sample with the smallest number of reads, and requiring a gene to be past this threshold in at least seven individuals - representing the number of individuals in the smallest sampling group. Genes were considered differentially expressed at a false discovery rate cut-off of 0.01 and a minimum four-fold difference in expression.

**Figure 1.**
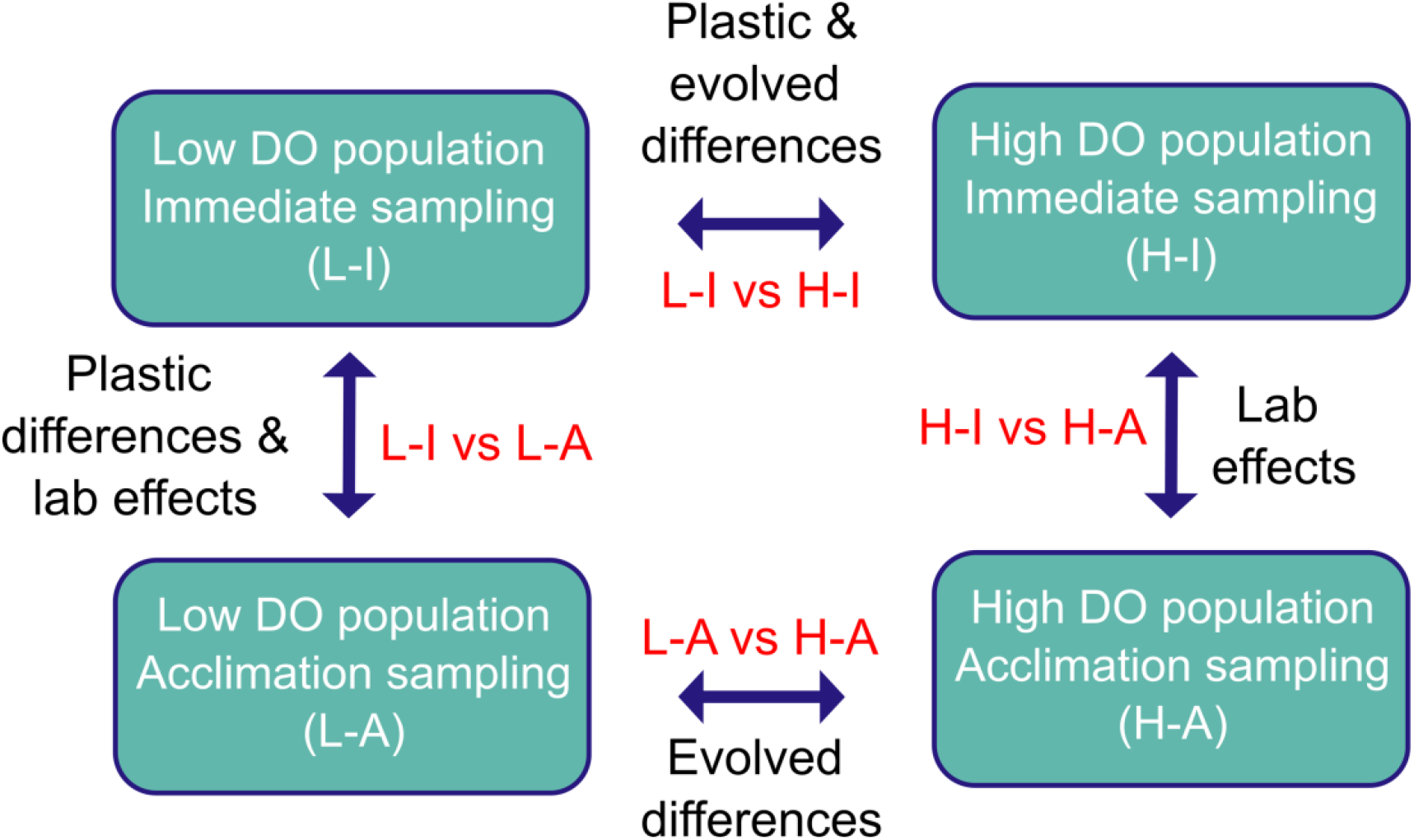
Pairwise comparisons made within each species for differential gene expression analysis. DO = dissolved oxygen.

**Figure 2.**
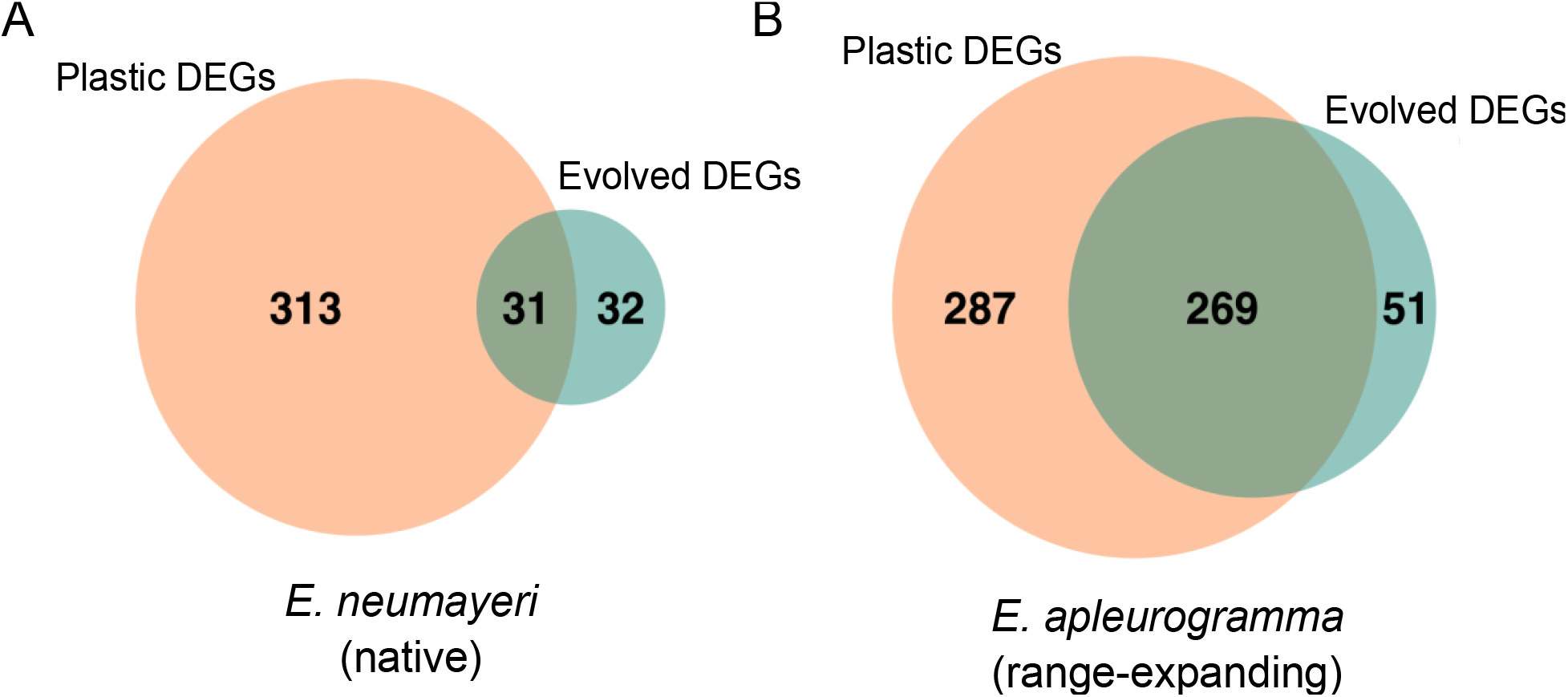
Venn diagram of plastic and evolved differentially expressed genes (DEGs) and the overlap between the two for (A) *E. neumayeri* (native) and (B) *E. apleurogramma* (range-expanding). Evolved DEGs were identified through the high dissolved oxygen (H-DO) population, acclimation sampling vs low dissolved oxygen population (L-DO), acclimation sampling comparison. Plastic DEGs were identified by finding the DEGs from the L-DO, immediate sampling vs L-DO, acclimation sampling comparison and then excluding DEGs that were also found in the H-DO, immediate sampling vs H-DO, acclimation sampling comparison.

### Cluster analyses

Further statistical analysis was performed on trimmed mean of M values (TMM) normalized, log2-transformed, and median centered gene expression values. To evaluate trends in total gene expression and differential gene expression, we ran principal component analyses (PCA) on all genes and on all differentially expressed genes. We assessed the PCAs for effects of holding pond, population (low- and high-DO), and sample type (immediate vs acclimation). We also ran hierarchical clustering with Euclidean distance and Ward’s linkage on samples using *cluster* v2.1.4 (Maechler et al., 2022).

To identify and visualize expression patterns across genes, we used *MFuzz* v2.60.0 (Kumar & Futschik, 2007) to perform soft (fuzzy c-means) clustering on our differentially expressed genes. This package groups genes with similar expression patterns together and assigns each gene a membership value to the cluster it is assigned to, representing how closely its expression aligns with the rest of the cluster. First, we estimated the optimal fuzzifier parameter using the *mestimate* function and then used *Dmin* and *cselection* to investigate the potential optimal number of clusters. For both species, the suggested number of clusters was 2 with the minimum centroid distance rapidly decreasing after 16 clusters. However, it is advised to visually review the data before choosing the number of optimal clusters as these tools may not always accurately identify all patterns, so we performed repeated clustering for a range of cluster numbers (c = 2-20) and visually assessed expression patterns to determine the number of clusters at which no uniquely shaped expression patterns were collapsed. For *E. neumayeri*, at the suggested number of clusters (c = 2) many unique expression profiles were collapsed which then became separate at c = 9; but at c > 9 redundant expression patterns became apparent. Therefore, we selected c = 9. Following the same reasoning, we selected c = 7 for *E. apleurogramma*. After selecting the final cluster number, we visualized expression across these clusters, requiring a minimum membership value of 0.7 for all genes. Lastly, we constructed a heatmap that displayed the clustering results for samples and DEGs with relative gene expression using *pheatmap* v1.0.12 (https://CRAN.R-project.org/package=pheatmap) and visually identified clusters of interest that contained genes with expression patterns showing differential expression between H-I samples and L-I samples that are no longer differentially expressed between the H-A vs L-A samples.

### Comparing plastic to evolutionary changes in gene expression within and between species

We identified three sets of genes: 1. Evolved changes: identified in the H-A vs L-A comparison; 2. Plastic changes: identified as DEGs in the L-I vs L-A comparison (lab effects and plasticity) that are not present in the H-I vs H-A comparison (lab effects) and, 3. Shared changes: identified by finding the overlap between the evolved DEGs and plastic DEGs. We compared log2-fold change (FC) in the shared DEGs using Pearson’s correlation to determine if these DEGs showed shifts in gene expression in the same or opposite direction. If a gene was upregulated or downregulated in both the L-A samples and the H-A samples, it was said to be in the same direction. We compared the observed correlation to a distribution produced through permutation by randomly sampling the number of shared genes (31 for *E. neumayeri* and 269 for *E. apleurogramma*) from all genes retained in the DEG analysis 10,000 times and recalculating Pearson’s correlation. We also used a Chi-square test with Yates correction to determine if there was a higher or lower proportion of evolutionary divergence DEGs overlapping with the plastic genes than expected. Within species, we ran a Chi-square test with Yates’ correction to test for differences in the proportion of significant DEGs for evolved and plastic DEG gene sets. To compare the plastic responses and evolved divergence between species, we ran Mann-Whitney U tests on the average magnitude of log2-FC in DEGs from the L-A vs L-I and the H-A. vs L-A comparisons between species. We used the *rstatix* package to calculate the effect sizes of the comparisons (Kassambara, 2023).

### Gene ontology enrichment analysis

We conducted gene ontology enrichment analysis on the gene clusters of interest identified in the cluster analysis using the package *Goseq* v3.17 (Young et al., 2010) to perform gene ontology (GO) enrichment analysis while correcting for gene length bias. All three GO branches (Cellular Components, Biological Processes, and Molecular Functions) were used to test for enrichment using the Wallenius approximation while restricting to a background list of the genes that were retained for the differential expression analysis after filtering for minimum expression. Then, we corrected *p*-values for multiple testing by converting to *q*-values using the package *qvalue* v2.32.0 (Storey et al., 2023) and considered terms as significantly enriched at a false discovery rate of *q* < 0.05. To further analyze and plot enriched GO terms we used REVIGO (http://revigo.irb.hr/) on all significant GO terms in the biological process category using default settings (whole UniProt database, medium list size, and SimRel similarity measure). REVIGO removes redundant GO terms and performs SimRel clustering to plot the similarity of given GO terms in semantic space (Supek et al., 2011).

## Results

### Sequence count overview, Trinity assemblies, and differential gene expression analysis

*E. neumayeri* (EN) samples had a sequencing depth range of 11.6 million to 39.4 million trimmed paired end (PE) reads (average 20,929,949 +/- 5,964,781) and *E. apleurogramma* (EA) samples had a range of 12.4 million to 71.6 million reads (average 33,856,401 +/- 14,498,298) (Supplemental Table 2 and 3). The Trinity assembly generated for *E. neumayeri* contained 1,094,565 transcript contigs grouped into 611,156 “genes” (median contig length: 374, N50 of 1009) (Supplemental Table 4). The *E. apleurogramma* Trinity assembly had 899,947 transcript contigs grouped into 546,319 “genes” (median contig length: 406, N50 of 1197) (Supplemental Table 4). BUSCO reports generated for each assembly indicated near complete gene sequence information for 91.6% of genes for *E. neumayeri* and 92.7% of genes for *E. apleurogramma* (Table 2). An average of 97% and 98.4% of reads per sample aligned back to the *E. neumayeri* and *E. apleurogramma* assemblies respectively with most of these reads mapped as proper pairs (Supplemental Tables 2 and 3). Using Trinotate to annotate the *E. neumayeri* assembly, we found 185,053 transcripts matching 35,511 unique Swiss-Prot proteins, 11,869 of which matched at least 80% of the protein’s length. For *E. apleurogramma*, we found 181,074 transcripts matching 34,996 unique Swiss-Prot proteins with 12,086 of which matched at least 80% of the protein’s length. After filtering, 33,776 and 73,898 Trinity “genes” were retained for differential gene expression analysis for *E. apleurogramma* and *E. neumayeri* respectively. We identified a total of 1,233 differentially expressed genes (DEGs) across 997 unique genes for *E. neumayeri* (see Supplemental Figure 1 for full breakdown).

**Table 2.** BUSCO reports for each species.

| Species | Summary in BUSCO annotation |
| --- | --- |
| *E. neumayeri* | C:91.6% [S:18.5%,D73.1%],F:4.8%,M:3.6%,n:3354] |
| *E. apleurogramma* | C:92.7% [S:15.9%,D76.8%],F:4.7%,M:2.6%,n:3354] |

### Cluster analyses

For the PCA on all genes in *E. neumayeri*, there was divergence between population (low-and high-DO) and sample type (immediate vs acclimation), although there was some overlap between the acclimation samples from both populations (Figure 3A). In *E. apleurogramma*, the PCA on all genes showed high levels of overlap between the immediate samples from each population, whereas the acclimation samples from each population showed divergence along PC1, with the L-A samples diverging the most from the immediate samples (Figure 3B). In the PCA on DEGs, there was clear clustering by population and sample type for both species (Figure 3C and D). *E. neumayeri* showed slight overlap between the acclimation samples (Figure 3C),while *E. apleurogramma* showed separation between those samples and instead slight overlap between the immediate samples (Figure 3D). When analyzing the PCA for any impact of acclimation pond on sample clustering, we found that samples did not cluster by acclimation pond suggesting no or little impact (Supplemental Figure 2). The hierarchical clustering on samples showed H-I and L-I samples clustering together for both species, however, for *E. apleurogramma* this cluster is then nested within the H-A samples and the L-A samples are the least like the rest (Figure 4A and B). For *E. neumayeri*, the H-A and L-A samples form a separate cluster (Figure 4A). Fuzzy cluster analysis on the DEGs using soft clustering identified 9 clusters for *E. neumayeri* and 7 clusters for *E.* apleurogramma (Supplemental Figures 3 and 4). Of these clusters, cluster 5 was determined to be of interest and potentially involved in plastic responses to DO levels in *E. apleurogramma* (Figure 4B) and cluster 5 and 6 were of interest for *E. neumayeri* (Figure 4A).

**Figure 3.**
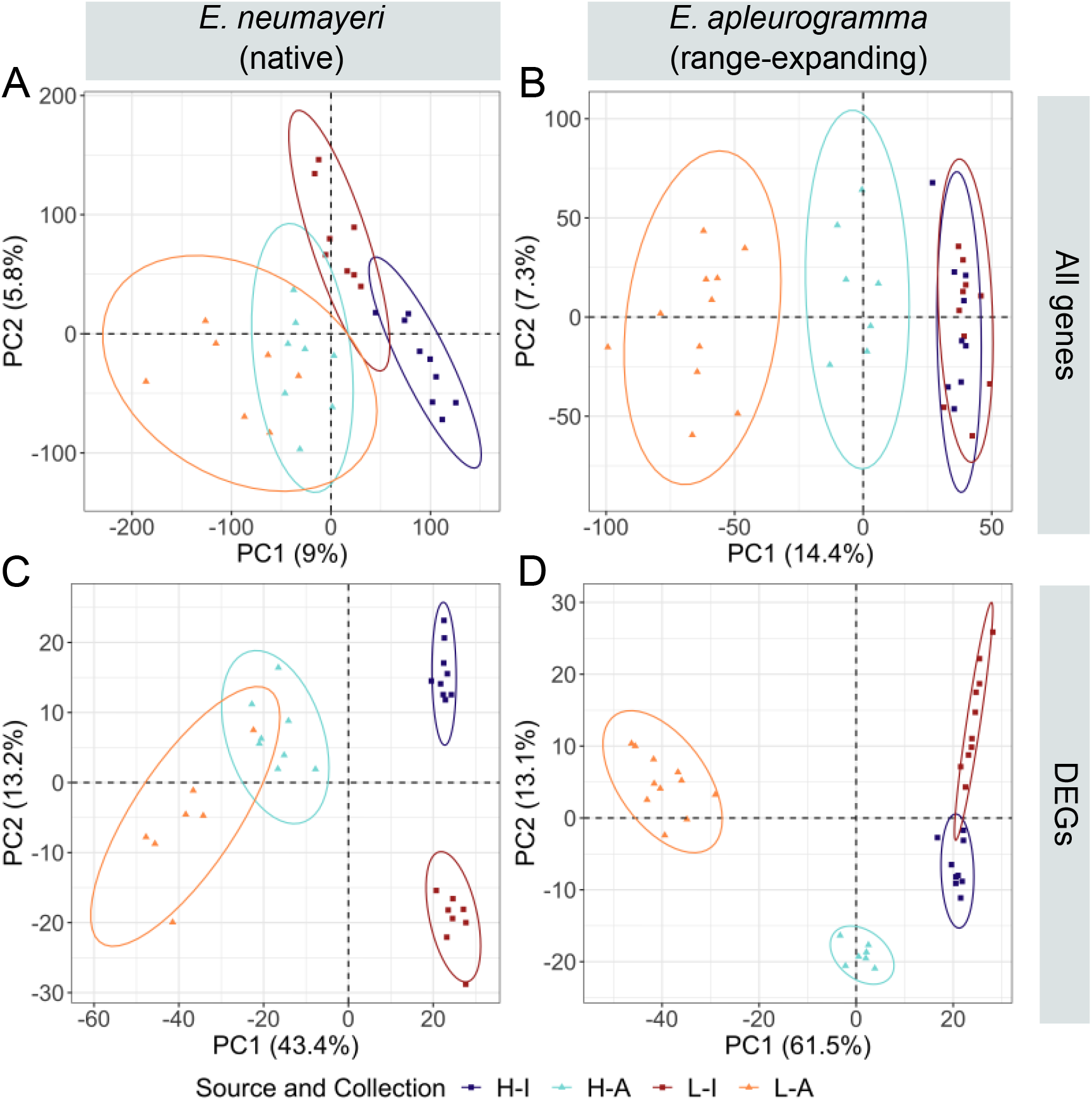
Principle component analysis (PCA) on all expression (A and B) and all differentially expressed genes (DEGs) (C and D) in *E. neumayeri* (native) (A and C) and *E. apleurogramma* (range-expanding) (B and D). Gene expression values were trimmed mean of M values (TMM) normalized, log2 transformed, and median centered prior to analysis. L = low DO population, H = high DO population, I = immediately sampled, A = sampled after acclimation.

**Figure 4.**
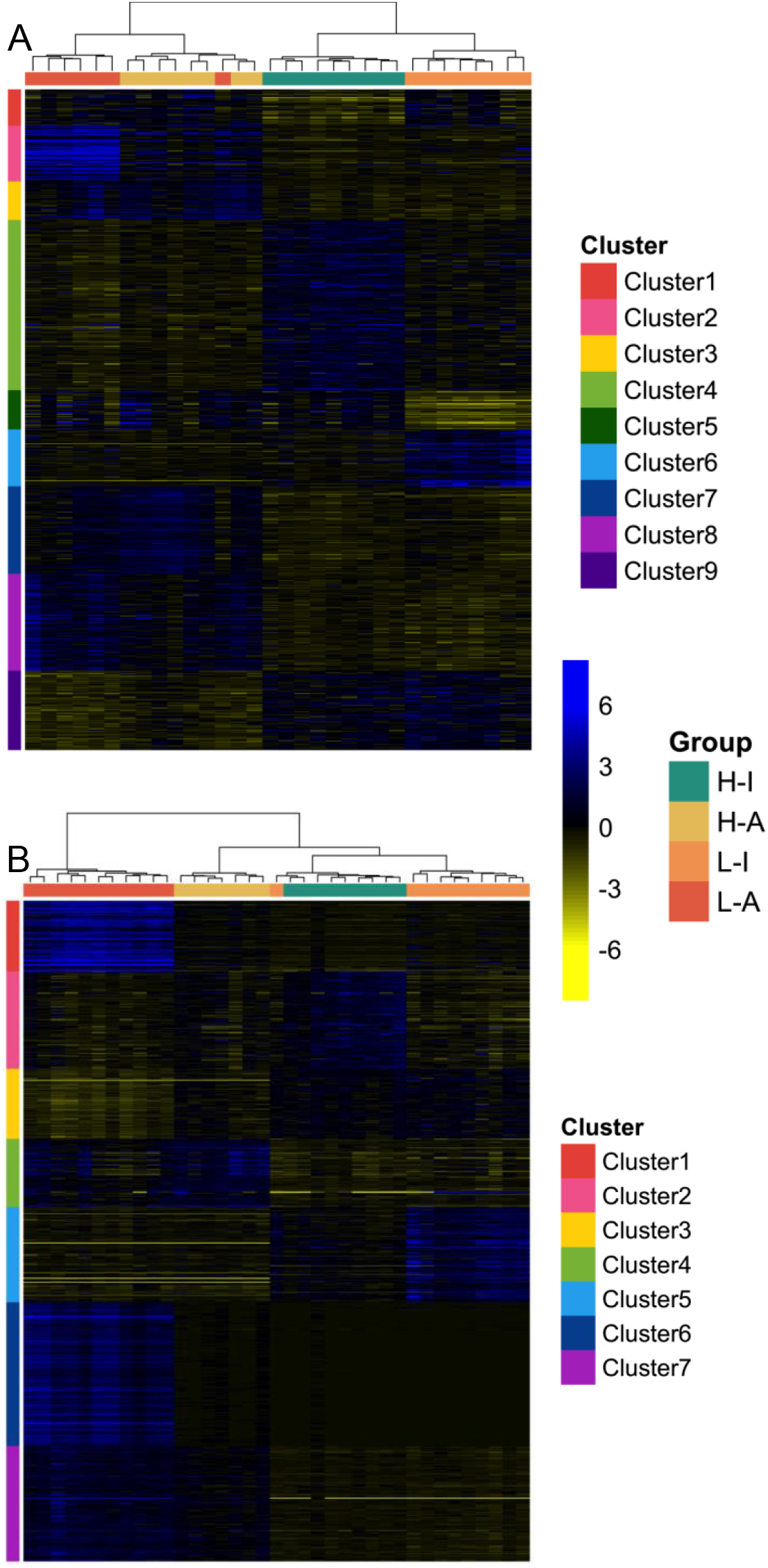
Heatmaps of differentially expressed genes (DEGs) with cluster analysis for (A) *E. neumayeri* (native) and (B) *E. apleurogramma* (range-expanding). Gene expression values are trimmed mean of M values (TMM) normalized, log2 transformed, and median centered. Hierarchical clustering with Euclidean distance and Ward’s linkage was run on samples and results are shown along the top of the heatmap. Soft (fuzzy c-means) clustering was run on DEGs to identify expression patterns with clustering results shown along the side of the heatmap. Clusters of interest that contained genes with expression patterns showing differential expression between H-I samples and L-I. samples that are no longer differentially expressed between the L-A and H-A samples were clusters 5 and 6 for *E. neumayeri* and cluster 5 for *E. apleurogramma*. L = low DO population, H = high DO population, I = immediately sampled, A = sampled after acclimation.

### Comparison between plastic and evolved gene expression

We identified 344 plastic, 63 evolved, and 31 shared DEGs for *E. neumayeri* (Figure 2A). *E. apleurogramma* had 556 plastic, 320 evolved, and 269 shared DEGs (Figure 2B). Log2-FC was highly negatively correlated between shared DEGs in both species (*E. neumayeri*: Pearson’s correlation = -0.838; *E. apleurogramma*: Pearson’s correlation = -0.792) (Figure 5A and B). We compared this result to a permutation test where we randomly sampled the number of DEGs retained in the analysis out of all genes 10,000 times and found that the observed correlation was stronger than expected by chance for both species (*p* = 0.0458 for *E. neumayeri* and *p* < 0.0001 for *E. apleurogramma*) (Figure 5A and B). There were no shared DEGs that showed changes in the same direction. In addition, genes showing evolutionary divergence overlapped with those showing plastic divergence at a higher rate (EN: 9.01%, EA: 48.38%) than they did within all genes (EN: 0.12%, EA: 1.05%) for both species (EN: X^2^ = 1537.6, df = 1, *p* < 0.0001; EA: X^2^ = 6560, df = 1, *p* < 0.0001).

**Figure 5.**
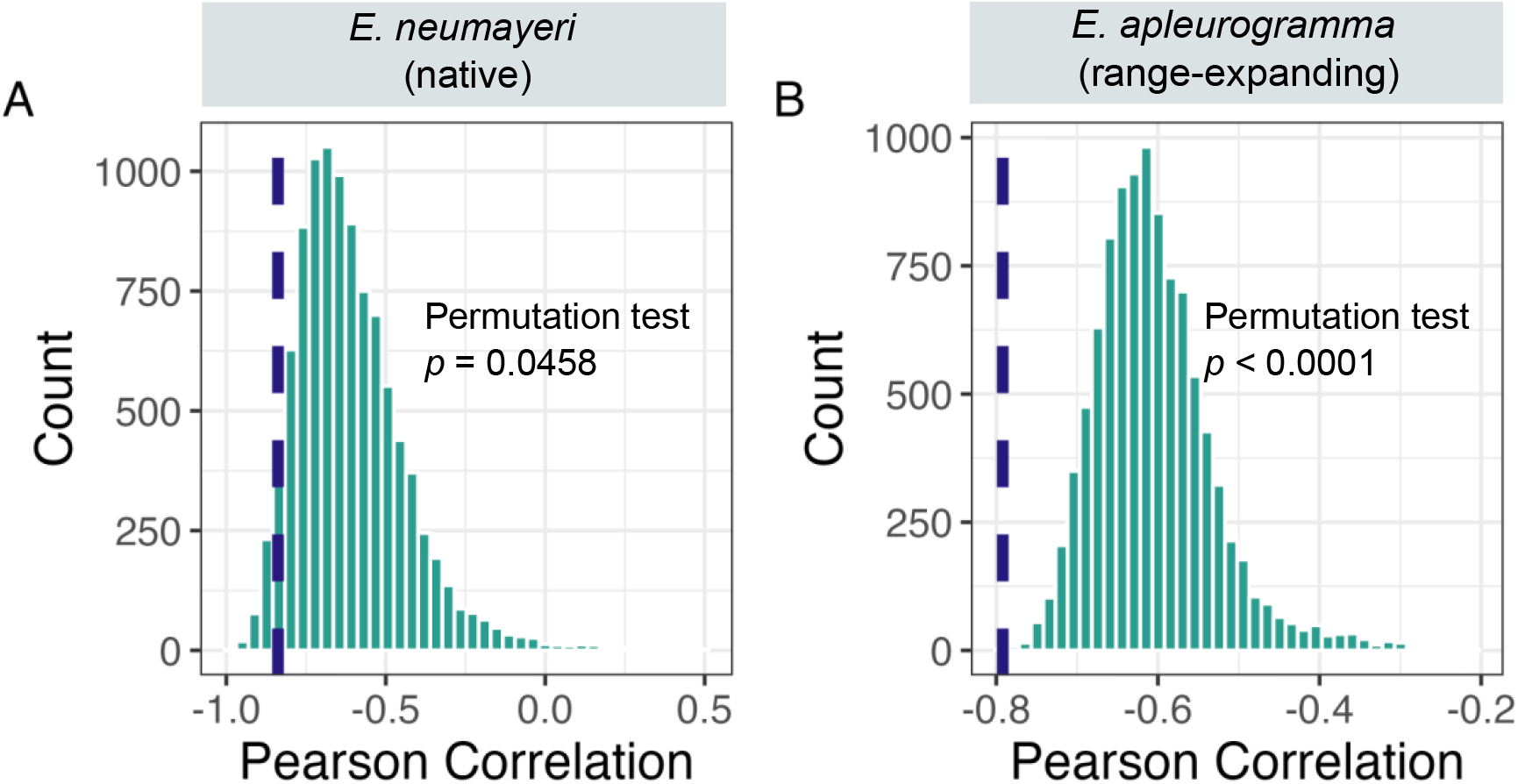
Correlation between log2 fold change in the evolutionary divergence and plastic shifts in (A) *E. neumayeri* and (B) *E. apleurogramma*. We ran permutation tests where we randomly sampled the number of DEGs that overlap out of all genes 10,000 times for each species and then recalculated Pearson’s correlation. The distribution of correlations is displayed along with the observed correlation indicated by the dashed line.

Both species showed a higher proportion of significant plastic DEGs than evolved DEGs (EN: X^2^ = 230.5, df = 1, *p* < 0.0001; EA: X^2^ = 58.5, df = 1, *p* < 0.0001). *E. apleurogramma* had a higher median and larger interquartile range (IQR) of magnitude log2-FC than *E. neumayeri* in plastic DEGs (EA: median = 4.05, IQR = 7.14; EN: median = 2.52, IQR = 1.11). In evolved DEGs, *E. apleurogramma* had a higher median but a smaller IQR of magnitude log2 FC than *E.* neumayeri (*E. apleurogramma*: median = 7.30, IQR = 2.48; *E. neumayeri*: median = 5.31, IQR = 3.91). These differences were significant in the Mann-Whitney U tests (*p* < 0.0001 for both comparisons; Figure 6). However, the effect size for the comparison between the plastic DEGs was smaller (r = 0.28) than that of the evolved DEGs (r = 0.318).

**Figure 6.**
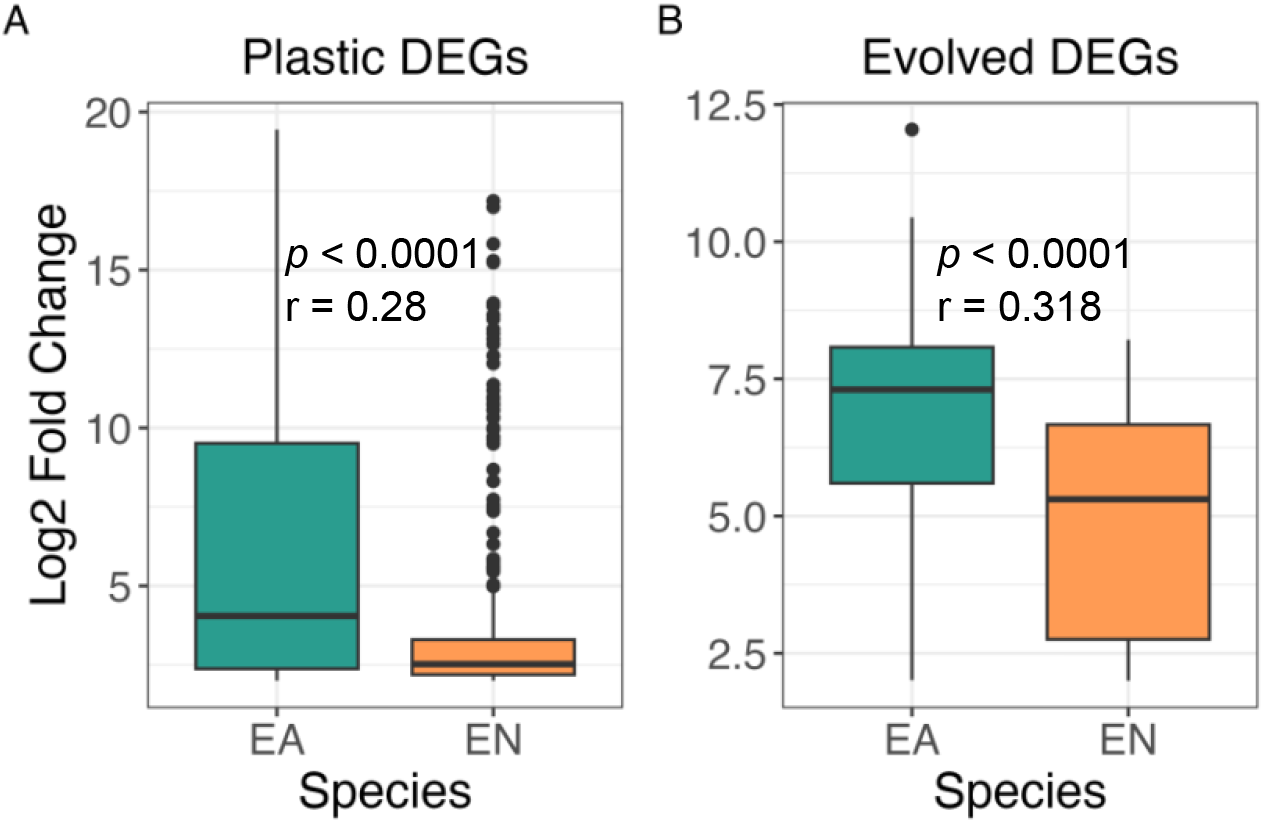
Difference in average magnitude log2 fold change between species for (A) Plastic changes, or (B) evolutionary changes. Evolved DEGs were identified through the high dissolved oxygen (H-DO) population, acclimation sampling vs low dissolved oxygen population (L-DO), acclimation sampling comparison. Plastic DEGs were identified by finding the DEGs from the L-DO, immediate sampling vs L-DO, acclimation sampling comparison and then excluding DEGs that were also found in the H-DO, immediate sampling vs H-DO, acclimation sampling comparison. Significance was tested using Mann-Whitney U tests. EA = *E. apleurogramma* (range-expanding), EN = *E. neumayeri* (native).

### Gene ontology enrichment analysis and REVIGO

For *E. neumayeri*, we conducted GO enrichment analysis and then REVIGO on clusters 5 and 6. Cluster 5 contained genes with decreased expression in the L-I samples compared to the rest of the samples (Figure 4 A). This cluster showed significant GO terms related to immune responses and regulation of the immune system (Supplemental Figure 5 A). Cluster 6 contained genes that were upregulated in L-I samples (Figure 4 A) and this cluster contained significant GO terms related to cellular responses to hypoxia, protein hydroxylation, and metabolic processes (Supplemental Figure 5 B). *E. apleurogramma* had GO enrichment analysis and REVIGO run on cluster 5 which contained genes that were upregulated in the L-I samples (Figure 4 B) and showed GO terms related to responses to hypoxia and nitric oxide, protein hydroxylation, and postsynaptic processes (Supplemental Figure 5 C; Supplemental Tables 5 and 6 for all significant GO terms).

## Discussion

We compared gene expression plasticity in response to DO levels between two species that have experienced different timescales of exposure to a naturally varying environment: one range-expanding species (*E. apleurogramma*) and one native species (*E. neumayeri*). Sampling fish from low-DO and high-DO field populations both immediately after capture and after a high-DO acclimation trial allowed us to disentangle evolutionary differences from short-term plastic differences in each species. Across our analyses, we found results that point to the importance of maladaptive plasticity in promoting divergence between high- and low-DO populations through counter-gradient variation for both species. However, our results suggest that the counter-gradient variation may be stronger in the recently colonizing species and could be facilitating colonization by promoting rapid genetic divergence between low-DO and high-DO populations. We also identified gene clusters involved in responses to DO levels with significant GO terms in two non-model fish species, many of which match findings found in mammalian study systems.

### Samples clustered differently by population origin and DO exposure for each species

Cluster analyses showed higher than expected levels of similarity between the field populations in both species. In the PCA on all genes, there is complete overlap of the high-DO and low-DO immediate samples for *E. apleurogramma* (range-expanding), while in *E. neumayeri* (native) there is slight overlap between the immediate samples but much more overlap between the acclimation samples. The PCA on DEGs indicate a similar pattern, with more overlap between the immediate samples of each population for *E. apleurogramma* than *E. neumayeri*. It is unknown which DO environment the colonizing *E. apleurogramma* individuals originated from, but it is expected that they primarily come from the high-DO population because this population has a much more direct route for migration to the RSS. Additionally, the field populations of *E. apleurogramma* have had much less time to diverge from each other than the field populations of *E. neumayeri*. Therefore, we would expect to see closer clustering between the immediate samples in *E. apleurogramma* than *E. neumayeri*. In the hierarchical analysis of all DEGs, the immediate samples clustered most closely together in both species. This is expected for *E. apleurogramma* due to the reasons outlined above but is surprising for *E. neumayeri*. The clustering of the immediate samples could be due to the laboratory conditions differing from the field (differences in water quality, food availability, etc.) which may have an impact on gene expression such that there is clustering within sampling type (acclimation vs immediate). Alternatively, this pattern could be due to the observed counter-gradient variation (see below).

### Counter-gradient variation in DEGs that overlap between plastic and evolved changes

We identified evolved DEGs (H-A vs the L-A) and plastic DEGs (L-I vs L-A minus H-I vs H-A). By comparing these sets of DEGs, we found that the majority did not overlap, suggesting that plasticity and evolved divergence occurs mostly in different subsets of genes. This finding is consistent with other studies. For example, in killifish (*Fundulus heteroclitus*) that experience varying temperatures there was very little overlap between plasticity DEGs and adaptation DEGs (Dayan et al., 2015). This result may suggest that plasticity impedes adaptive divergence since the genes that experience plasticity do not diverge between populations. One way to determine if plasticity is impeding evolutionary divergence is to compare the proportion of evolutionary divergence DEGs in all genes to the proportion in plastic genes. We found that there was a significantly higher proportion of evolutionary divergence genes within the plastic gene set than within all genes for both species, suggesting that plasticity does not impede genetic divergence. The subset of genes that did overlap, demonstrating both plastic and evolved differences, were highly negatively correlated in both species. This correlation was stronger than expected by chance as tested in a permutation analyses. Additionally, the fold change of expression was found to operate in the opposite direction for all shared DEGs. Adaptive differences that occur in the opposite direction as the plastic differences are suggestive of counter-gradient variation (Conover & Schultz, 1995).

Counter-gradient variation can evolve through genetic compensation, a subset of genetic accommodation, where a plastic change in phenotype reduces fitness in a new environment but selection subsequently acts to shift the phenotype back to the ancestral state without reducing phenotypic plasticity (Grether, 2005). As a result, genetic compensation may lead to populations from different environments displaying higher trait similarity in the field than when acclimated in a common environment. In the cluster analysis, the range-expanding *E. apleurogramma* showed more similarity between populations from different DO habitats when they were sampled immediately in the field than after they had been acclimated to normoxia. This pattern may be due to genetic compensation acting to reduce phenotypic variation between the populations. However, this pattern was not observed in the native *E. neumayeri,* which may suggest that this species is experiencing less counter-gradient variation. This could be due to *E. neumayeri* expressing less maladaptive plasticity than *E. apleurogramma*. Indeed, *E. neumayeri* showed consistently fewer counter-gradient genes than *E. apleurogramma*. However, it is unclear whether *E. neumayeri* has always possessed fewer counter-gradient genes or if there might have been similar levels of counter-gradient variation during initial stages of colonization which were then reduced over time. In guppies (*Poecilia reticulata*), between two lineages that have shown parallel evolution in response to high predation, the lineage that was more recently diverged showed a stronger signature of nonadaptive plasticity than the older lineage (Fischer et al., 2021). This suggests that studies must consider how plasticity impacts divergence across all stages of colonization to fully understand its role.

Previous studies have found conflicting results for counter-gradient variation in gene expression in fish. One study on guppies adapting to predator free environments found that 89% of transcripts showed shifts in gene expression that were in the opposite direction of evolved changes and concluded that maladaptive plasticity potentiates the rapid evolution of brain gene expression during the early stages of adaptation (Ghalambor et al., 2015). However, this study was criticized for making conclusions based on gene expression data only and collecting no data on organismal plasticity directly (van Gestel & Weissing, 2018). It is important to consider that gene expression is only one measurement of plasticity, and the complexity of regulation mechanisms make it possible for divergent changes in gene expression to lead to convergent phenotypes. Indeed, another study investigating gene expression directly from the proteome in populations of European grayling (*Thymallus thymallus*) instead found that plastic and evolved changes were in the same direction (Mäkinen et al., 2016). Further, a follow up study on guppies again showed nonadaptive plasticity but also suggested that alternative transcriptional configurations could be associated with shared phenotypes across distinct evolutionary lineages (Fischer et al., 2021). Therefore, our results are consistent with counter-gradient variation, however, further studies are needed to fully elucidate the role of maladaptive plasticity in this system.

### Higher plasticity and evolutionary divergence in E. apleurogramma

One way to assess the relative levels of plastic and evolved divergence is to compare the proportion of genes that are significantly differentially expressed for each type of change. We found that there was a significantly higher proportion of plastic DEGs than evolved DEGs for both species. This result is expected for *E. apleurogramma* which has had less time for populations in low- and high-DO habitats to show evolved divergence, however, it is surprising for *E. neumayeri* which is expected to be genetically divergent between low- and high-DO sites. A level of DO induced plasticity is likely maintained in *E. neumayeri* despite evolved divergence across DO habitats since DO fluctuates seasonally (Chapman et al., 1999) and the proximity of the sites (∼200 m) means that some individuals may occasionally cross DO boundaries.

Further support for plasticity playing a more important role in *E. apleurogramma* than *E. neumayeri* comes from comparisons of log2-fold change (FC), which show that *E. apleurogramma* had a higher median magnitude of log2-FC for plastic DEGs. Interestingly, the range of magnitude log2-FC for the plastic DEGs for *E. apleurogramma* was much larger than the range for *E. neumayeri*. Although, *E. apleurogramma* has only a slightly larger magnitude of plastic change than *E. neumayeri*, it possesses some genes that show very strong plastic responses. If these highly plastic genes are adaptive, they may be responsible for facilitating *E. apleurogramma’s* coloniaztion of the RSS by enabling the species to persist in the low-DO environments and thereby allowing for genetic assimilation (Crispo, 2007; Schlichting & Wund, 2014). However, the counter-gradient variation we observed indicates that these plastic genes could also be maladaptive. Maladaptive plasticity has been hypothesized to aid adaptive divergence in some cases by increasing the strength of selection (Ghalambor et al., 2007). This could facilitate colonization by increasing the speed of genetic adaptation. Experimental range shifts of the seed beetle, *Callosobruchus macuulatus*, into cooler and more variable conditions showed that heat and cold tolerance rapidly evolved, however, this adaptation was associated with maladaptive plasticity in the novel conditions which resulted in a pattern of counter-gradient variation (Leonard & Lancaster, 2020). Beetles that colonized only colder but not more variable environments expressed only adaptive plasticity and no evolved response. The RSS has temporal and spatial variation in DO levels (Chapman et al., 1999) that may be promoting rapid adaptation through maladaptive plasticity and counter-gradient variation. Multiple studies have found evidence that swamp populations of *E. neumayeri* may be maladapted to their environment. One study found swamp populations have lower fecundity, reproductive investment, and condition (Baltazar, 2015) while another showed no growth advantage for swamp fish over stream fish in a swamp environment (Martínez et al., 2011). Our results suggest these patterns could be due to maladaptive plasticity in gene expression.

We also found that *E. apleurogramma* had a higher median magnitude of log2-FC for evolved DEGs relative to *E. neumayeri*. This is surprising given that *E. neumayeri* has a longer evolutionary history in the RSS and was expected to be more divergent between high- and low-DO populations than *E. apleurogramma* which is new to the RSS. This finding could suggest that there are more migrants exchanged between populations of *E. neumayeri* than previously hypothesized. One mark recapture study on *E. neumayeri* found that 7% of individuals dispersed from their location of capture with some individuals travelling across DO environments (Chapman et al., 1999). Indeed, in the cluster analyses there are a few instances where individual samples group more closely within the other population. This finding may also give further support to the hypothesis that the observed counter-gradient variation could be contributing to the development of rapid genetic divergence between the two populations of *E. apleurogramma*.

In contrast, *E. neumayeri* shows smaller variation in plasticity between genes and a lower magnitude log2-FC for plasticity and evolutionary divergence. It is hypothesized that costs of plasticity lead to decreasing levels of plasticity over time as evolutionary changes begin to take effect (Crispo, 2007). The smaller range of plasticity seen in *E. neumayeri* could suggest that the level of plasticity has been reduced by selection. The lower magnitude of plasticity could alternatively suggest that the counter-gradient variation was less strong in this species, which could explain the smaller evolutionary divergence between the two populations. In tree sparrows (*Passer montanus*), the amount of genetic divergence between populations experiencing varying oxygen environments due to altitude depends on the magnitude of counter-gradient variation (She et al., 2023). As previously mentioned, cluster results suggest that *E. apleurogramma* may be experiencing stronger counter-gradient variation than *E. neumayeri,* which could result in more genetic divergence between populations.

### Gene clustering and identification of genes related to hypoxia responses

Using soft clustering, we identified two clusters of interest for *E. neumayeri* and one cluster of interest for *E. apleurogramma* that had expression profiles suggesting involvement in plastic responses to DO levels. The gene ontology analysis of these clusters of interest identified genes involved in responses to hypoxia for both species that are upregulated in the low-DO, immediate samples. In mammals, research on hypoxia has identified the hypoxia-inducible factor (HIF) that regulates gene expression cascades in response to lower oxygen levels (Nikinmaa & Rees, 2005). HIF mediated gene expression is oxygen sensitive due in part to the degradation of the HIF-α subunit that is mediated by an oxygen-dependent degradation (ODD) domain. In this domain, specific proline residues are hydroxylated and then degraded under normoxic conditions. Under hypoxic conditions, hydroxylation does not occur and HIF-α accumulates and then binds to promoter or enhancer regions of hypoxia-inducible genes. Interestingly, there was upregulation of genes involved in protein hydroxylation or proline hydroxylation and in genes related to 4-hydroxyproline metabolic processes in both species. This may be evidence that these species utilize different oxygen dependent steps in HIF gene expression pathways. Alternatively, these shifts in expression may represent mechanisms to reduce the impact of hypoxia. Previous studies have found that HIF pathways are less activated in human populations that are adapted to high-elevation compared to populations at sea level (Storz, 2021). Therefore, it is possible that the fish show adaptation to low-DO that allows for the suppression of HIF pathways. Other gene groups known to be involved in HIF gene expression cascades were found to be significantly enriched as well. In *E. apleurogramma*, genes related to the regulation of CAMKK-AMPK signalling cascade were upregulated under low-DO which has been previously found to be upregulated in gill tissue under hypoxic stress (Ren et al., 2022). There was also an increase in expression of genes related to responses to nitric oxide, which mediates vasodilation to help deliver more oxygen to tissues (Ho et al., 2012).

In *E. neumayeri*, we also identified a cluster containing genes that were down-regulated in L-I samples. Most of the significant GO terms in this cluster were related to immune and defense responses. Down regulation of immune related genes under hypoxia stress has also been found in zebrafish (*Danio rerio*) (van der Meer et al., 2005), tilapia (*Oreochromis niloticus*) (Li et al., 2017), and large yellow croaker (*Larimichthys crocea*) (Mu et al., 2020) in various tissues including in gill tissue which could be especially detrimental to the health of fish experiencing hypoxic conditions due to gill tissue being a primary barrier to pathogens.

### Future Directions

This study adds to the growing evidence that counter-gradient variation in gene expression plays a role in the early stages of colonization, however, there are several important issues that should be addressed in future research. One limitation is that we are assuming that the gene expression patterns displayed between the H-A vs the L-A comparison represent heritable differences between the two populations because they persist after the acclimation trial to a common DO environment. However, it is possible that this comparison also includes irreversible developmental plasticity. Future studies could disentangle levels of developmental plasticity from evolved divergence by raising multiple generations under acclimation trials. Additionally, work on developing analytical frameworks to quantify co-gradient and counter-gradient variation suggest that the best experimental design to decipher between the two is a reciprocal transplant design where individuals are exposed to both environments (Albecker et al., 2022). Low-DO acclimations are logistically difficult to run at field stations, however, to confirm whether there is indeed counter-gradient variation in these species, a future study should run the acclimation study in both low-DO and high-DO and try to apply these new analytical techniques. As previously mentioned, measuring plasticity at the phenotypic level would also further distinguish counter-gradient from co-gradient variation by determining which changes in gene expression result in divergent phenotypes and what the adaptive consequences are. Another limitation is that the acclimation used in this study represents a shift in only one environmental parameter whereas the low-DO and high-DO environments likely vary in many biotic and abiotic conditions that could covary with DO. While this study focuses on plastic and adaptive responses to DO, adaptation to these different environments likely requires plasticity or local adaptation in a suite of traits that may not be directly impacted by DO. For example, predation is often lower in low-DO environments, allowing for swamps to act as a refuge for prey species (Chapman et al., 2002). Future research could study multiple overlapping environmental parameters to obtain a more comprehensive understanding of the relationship between adaptive divergence and plasticity.

## Conclusion

In this study, we described gene expression responses to hypoxia in two fish species and compared plastic to evolved changes. We found that plastic changes mostly occur in different genes from evolutionary divergence and uncovered evidence suggesting counter-gradient variation in plasticity and evolved divergence in both a recently range-expanding and long-established species. This counter-gradient variation might be due to maladaptive plasticity that is being genetically compensated for. We suggest that plasticity may not need to be adaptive to facilitate colonization of novel environments; maladaptive plasticity may also aid colonization by increasing the strength of selection and promoting rapid adaptive genetic divergence. This study provides insight into how phenotypic plasticity and genetic divergence interact to shape populations diverging across varying environments.

## Acknowledgements

We would like to thank Allegra Pearce, Alexis Heckley, and members of the Barrett lab for providing comments on the manuscript. We thank Dr. Patrick Omeja and the field assistants of the Kibale Fish Project for assistance with field logistics.

JAF was supported by a Natural Sciences and Engineering Research Council (NSERC) Postgraduate scholarship – Doctoral (#PGSD3-559394-2021). RB was supported by an NSERC Discovery Grant #2019-04549 and a Canada Research Chair. The lab and field research for this project were also supported by an FRQNT grant FRQ-NT PR-298361 (RB, LJC) and an NSERC Discovery Grant #2020-04315 (LJC).

## Data Accessibility and Benefit Sharing

### Data accessibility statement

The data presented in the study will be deposited in the NCBI Sequence Read Archive (SRA) repository, accession number TBD.

### Author contributions

Janay Fox carried out laboratory work, analyzed data, and wrote the manuscript. David Hunt carried out fieldwork and gave feedback on the manuscript. Andrew Hendry, Lauren Chapman, and Rowan Barrett conceptualized the experimental design, provided funding, and gave feedback on the manuscript.

## Notes

### Competing Interest Statement

The authors have declared no competing interest.

