## Supplemental Information for "Counter-gradient variation in gene expression between fish populations facilitates colonization of low-dissolved oxygen environments"

### Table of Contents:


| | |
|---|---|
| **Supplemental Table 1. Temperature and dissolved oxygen (DO) data for acclimation ponds.** | Page 2 |
| **Supplemental Table 2. Sequencing depth before and after trimming and alignment rate for *E. neumayeri*.** | Page 2 |
| **Supplemental Table 3. Sequencing depth before and after trimming and alignment rate for *E. apluerogramma*.** | Page 3 |
| **Supplemental Table 4. Trinity assembly quality metrics.** | Page 5 |
| **Supplemental Table 5. Significant GO terms from gene ontology analysis on *E. neumayeri*.** | Page 5 |
| **Supplemental Table 6. Significant GO terms from gene ontology analysis on *E. neumayeri*.** | Page 13 |
| **Supplemental Figure 1. Number of differentially expressed genes identified for each type of comparison made for (A) *E. neumayeri* and (B) *E. apleurogramma*.** | Page 15 |
| **Supplemental Figure 1. Principle component analysis (PCA) on TMM normalized, log2 transformed, and median centered gene expression values for all genes and all differentially expressed genes in *E. neumayeri* and *E. apleurogramma* labelled with acclimation pond.** | Page 16 |
| **Supplemental Figure 2. Soft cluster results for *E. neumayeri*.** | Page 17 |
| **Supplemental Figure 3. Soft cluster results for *E. apleurogramma*.** | Page 18 |

**Supplemental Table 1. Temperature and dissolved oxygen (DO) data for acclimation ponds.**

| Date | Time | Pond | DO (mg/L) | Temperature (C°) |
|---|---|---|---|---|
| 2017-06-06 | 14:18 | 1 | 7.19 | 21 |
| | | 2 | 7.2 | 20.8 |
| 2017-06-07 | 08:36 | 1 | 6.23 | 18.6 |
| | | 2 | 6.07 | 18.6 |
| 2017-06-08 | 15:54 | 1 | 7.22 | 20 |
| | | 2 | 7.16 | 20 |
| 2017-06-09 | 08:49 | 1 | 7.24 | 18.4 |
| | | 2 | 7.28 | 18.4 |
| 2017-06-10 | 08:36 | 1 | 7.25 | 18.2 |
| | | 2 | 7.22 | 18.2 |
| 2017-06-10 | 18:57 | 1 | 7.01 | 19.8 |
| | | 2 | 7.22 | 19.7 |
| 2017-06-12 | 08:31 | 1 | 6.82 | 19.2 |
| | | 2 | 6.97 | 18.9 |
| 2017-06-13 | 08:05 | 1 | 7.16 | 19.2 |
| | | 2 | 7.24 | 19 |
| 2017-06-14 | 08:35 | 1 | 6.03 | 18.7 |
| | | 2 | 6.67 | 18.7 |
| 2017-06-16 | 08:59 | 1 | 5.63 | 17.6 |
| | | 2 | 5.93 | 17.7 |
| 2017-06-17 | 08:38 | 1 | 5.5 | 19 |
| | | 2 | 6.58 | 19 |
| 2017-06-19 | 08:38 | 1 | 6.28 | 18.1 |
| | | 2 | 6.72 | 18.1 |
| 2017-06-20 | 09:07 | 1 | 6.4 | 18 |
| | | 2 | 6.42 | 17.1 |

**Supplemental Table 2. Sequencing depth before and after trimming and alignment rate for *E. neumayeri*.**

| ID | Sample Type | Raw Reads | Trimmed Reads | Overall alignment rate | PE alignment rate |
|---|---|---|---|---|---|
| 0580g | High DO, Immediate | 30925396 | 20299416 | 78.03% | 54.22% |
| 0582g | High DO, Immediate | 59816531 | 39411990 | 97.81% | 78.88% |
| 0583g | High DO, Immediate | 25962009 | 17705552 | 97.70% | 78.58% |
| 0584g | High DO, Immediate | 51023628 | 33137424 | 98.03% | 80.21% |
| 0585g | High DO, Immediate | 33269240 | 19693986 | 97.60% | 80.46% |
| 0586g | High DO, Immediate | 43664226 | 26570548 | 97.44% | 80.87% |
| 0587g | High DO, Immediate | 30530817 | 18171247 | 98.13% | 81.35% |
| 0588g | High DO, Immediate | 38929518 | 24829617 | 97.97% | 80.02% |

| | | | | | |
|---|---|---|---|---|---|
| 0589g | High DO, Immediate | 39055528 | 26781325 | 97.91% | 79.49% |
| 0610g | Low DO, Immediate | 27447196 | 13113380 | 98.07% | 84.92% |
| 0611g | Low DO, Immediate | 37176685 | 23783078 | 97.79% | 81.39% |
| 0612g | Low DO, Immediate | 29927602 | 18271338 | 97.80% | 81.98% |
| 0613g | Low DO, Immediate | 31919349 | 19107021 | 97.43% | 80.07% |
| 0614g | Low DO, Immediate | 47856911 | 25960851 | 97.70% | 82.82% |
| 0615g | Low DO, Immediate | 39508854 | 23644457 | 97.57% | 81.18% |
| 0617g | Low DO, Acclimate | 20630851 | 11615166 | 97.54% | 82.76% |
| 0619g | Low DO, Acclimate | 35744996 | 22753475 | 96.65% | 78.81% |
| 0620g | Low DO, Acclimate | 45886182 | 31510431 | 97.60% | 80.28% |
| 0622g | Low DO, Acclimate | 25957989 | 17314191 | 97.51% | 80.14% |
| 0623g | High DO, Acclimate | 38726405 | 24888132 | 97.79% | 81.48% |
| 0624g | High DO, Acclimate | 31516742 | 18935983 | 97.76% | 82.84% |
| 0626g | High DO, Acclimate | 25707146 | 16260891 | 97.47% | 79.13% |
| 0627g | Low DO, Acclimate | 25602163 | 16618648 | 97.12% | 79.43% |
| 0628g | Low DO, Acclimate | 32611723 | 20398754 | 97.12% | 80.32% |
| 0630g | Low DO, Acclimate | 26322115 | 15819853 | 97.49% | 82.90% |
| 0637g | High DO, Acclimate | 29489321 | 22024505 | 97.81% | 80.96% |
| 0644g | Low DO, Acclimate | 22227435 | 12817512 | 97.34% | 80.11% |
| 0645g | High DO, Acclimate | 26910993 | 18552309 | 97.44% | 80.37% |
| 0646g | High DO, Acclimate | 23245963 | 14809341 | 97.52% | 80.66% |
| 0647g | Low DO, Acclimate | 30000379 | 17488329 | 97.56% | 82.86% |
| 0649g | High DO, Acclimate | 32501884 | 19528676 | 97.55% | 81.32% |
| 0653g | High DO, Acclimate | 28202553 | 17940936 | 97.53% | 80.39% |

PE = Paired end.

**Supplemental Table 3. Sequencing depth before and after trimming and alignment rate for *E. apluerogramma.***

| ID | Sample Type | Raw Reads | Trimmed Reads | Overall alignment rate | PE alignment rate |
|---|---|---|---|---|---|
| 0590g | High DO, Immediate | 45662428 | 27880512 | 98.74% | 84.38% |
| 0591g | High DO, Immediate | 57140662 | 39284731 | 98.33% | 80.80% |
| 0592g | High DO, Immediate | 83435957 | 57334959 | 98.52% | 82.64% |
| 0593g | High DO, Immediate | 40511172 | 26198344 | 98.23% | 82.08% |
| 0594g | High DO, Immediate | 34509610 | 22773622 | 98.14% | 79.58% |
| 0595g | High DO, Immediate | 113639460 | 71593578 | 98.46% | 83.43% |
| 0596g | High DO, Immediate | 29040114 | 19320044 | 98.36% | 81.99% |
| 0597g | High DO, Immediate | 51353039 | 32360208 | 98.72% | 84.40% |
| 0599g | High DO, Immediate | 84097603 | 52638277 | 98.56% | 83.22% |

| | | | | | |
|---|---|---|---|---|---|
| 0600g | Low DO, Immediate | 90858556 | 54028059 | 98.40% | 83.78% |
| 0601g | Low DO, Immediate | 43904740 | 29871954 | 98.56% | 82.21% |
| 0602g | Low DO, Immediate | 54912720 | 35947591 | 98.10% | 80.82% |
| 0603g | Low DO, Immediate | 51581862 | 31913253 | 98.72% | 84.73% |
| 0604g | Low DO, Immediate | 61695910 | 41022612 | 98.63% | 82.56% |
| 0605g | Low DO, Immediate | 24677521 | 14753556 | 98.30% | 83.54% |
| 0606g | Low DO, Immediate | 24059457 | 15768840 | 98.55% | 82.62% |
| 0607g | Low DO, Immediate | 103402281 | 65447207 | 98.06% | 81.35% |
| 0608g | Low DO, Immediate | 62045707 | 39073617 | 98.38% | 82.56% |
| 0609g | Low DO, Immediate | 27530898 | 17929150 | 98.50% | 82.33% |
| 0625g | Low DO, Acclimate | 69386988 | 46724552 | 98.24% | 81.11% |
| 0629g | Low DO, Acclimate | 42334128 | 31485949 | 98.04% | 77.67% |
| 0631g | High DO, Acclimate | 66757617 | 46791206 | 98.48% | 81.43% |
| 0632g | High DO, Acclimate | 25895959 | 15690372 | 98.75% | 84.22% |
| 0633g | High DO, Acclimate | 53252280 | 36477004 | 98.58% | 82.42% |
| 0634g | High DO, Acclimate | 51142795 | 35285543 | 98.73% | 82.43% |
| 0635g | Low DO, Acclimate | 32575677 | 20550292 | 98.43% | 82.33% |
| 0636g | High DO, Acclimate | 29489321 | 21350806 | 98.03% | 79.04% |
| 0638g | Low DO, Acclimate | 18525753 | 12421340 | 98.81% | 84.03% |
| 0639g | Low DO, Acclimate | 46351283 | 31836144 | 98.67% | 82.48% |
| 0640g | Low DO, Acclimate | 29667604 | 22042774 | 98.25% | 78.83% |
| 0641g | High DO, Acclimate | 84226972 | 52284477 | 98.53% | 83.70% |
| 0642g | Low DO, Acclimate | 48874205 | 34418379 | 98.25% | 80.03% |
| 0643g | Low DO, Acclimate | 43133678 | 27354030 | 98.20% | 81.53% |
| 0648g | Low DO, Acclimate | 40218298 | 25834891 | 98.24% | 81.97% |
| 0650g | Low DO, Acclimate | 53899556 | 31316480 | 98.39% | 83.70% |
| 0651g | Low DO, Acclimate | 79991012 | 48119914 | 98.59% | 84.01% |
| 0655g | High DO, Acclimate | 27640943 | 17562580 | 98.51% | 82.41% |

PE = Paired end.

**Supplemental Table 4. Trinity assembly quality metrics.**

| | Species | |
|---|---|---|
| *Nx Stats* | *E. neumayeri* | *E. apluerogramma* |
| N10 | 4397 | 4959 |
| N20 | 2760 | 3164 |
| N30 | 1904 | 2200 |
| N40 | 1383 | 1608 |

| | | |
|---|---|---|
| N50 | 1009 | 1197 |
| *Contig Length* | | |
| Median contig length | 374 | 406 |
| Average contig length | 671 | 750 |
| *Counts of transcripts* | | |
| Total Trinity transcripts | 1094565 | 899947 |
| Total Trinity "genes" | 611156 | 546319 |
| Percent GC | 41% | 41.2% |

Nx length statistics describe where at least x% of the assembled transcript nucleotides are found in contigs of at least Nx and are based on only the longest isoform per "gene".

**Supplemental Table 5. Significant GO terms from gene ontology analysis on *E. neumayeri*.**

| GO-term | Name | Ontology | *q*-value | Expression Pattern |
|---|---|---|---|---|
| GO:0001848 | complement binding | MF | <0.0001 | Cluster 5 |
| GO:0001872 | (1->3)-beta-D-glucan binding | MF | <0.0001 | Cluster 5 |
| GO:0002252 | immune effector process | BP | <0.0001 | Cluster 5 |
| GO:0005615 | extracellular space | CC | <0.0001 | Cluster 5 |
| GO:0006956 | complement activation | BP | <0.0001 | Cluster 5 |
| GO:0006957 | complement activation, alternative pathway | BP | <0.0001 | Cluster 5 |
| GO:0006958 | complement activation, classical pathway | BP | <0.0001 | Cluster 5 |
| GO:0006959 | humoral immune response | BP | <0.0001 | Cluster 5 |
| GO:0072376 | protein activation cascade | BP | <0.0001 | Cluster 5 |
| GO:0002253 | activation of immune response | BP | <0.0001 | Cluster 5 |
| GO:0030247 | polysaccharide binding | MF | <0.0001 | Cluster 5 |
| GO:0050778 | positive regulation of immune response | BP | <0.0001 | Cluster 5 |
| GO:0006955 | immune response | BP | <0.0001 | Cluster 5 |
| GO:0005576 | extracellular region | CC | <0.0001 | Cluster 5 |
| GO:0002376 | immune system process | BP | <0.0001 | Cluster 5 |
| GO:0050776 | regulation of immune response | BP | 0.0001 | Cluster 5 |
| GO:0003823 | antigen binding | MF | 0.0003 | Cluster 5 |
| GO:0045087 | innate immune response | BP | 0.0003 | Cluster 5 |
| GO:0004866 | endopeptidase inhibitor activity | MF | 0.0004 | Cluster 5 |
| GO:0030414 | peptidase inhibitor activity | MF | 0.0005 | Cluster 5 |
| GO:0061135 | endopeptidase regulator activity | MF | 0.0005 | Cluster 5 |
| GO:0006952 | defense response | BP | 0.0009 | Cluster 5 |
| GO:0002684 | positive regulation of immune system process | BP | 0.001 | Cluster 5 |
| GO:0009617 | response to bacterium | BP | 0.001 | Cluster 5 |

| | | | | |
|---|---|---|---|---|
| GO:0030449 | regulation of complement activation | BP | 0.002 | Cluster 5 |
| GO:2000257 | regulation of protein activation cascade | BP | 0.003 | Cluster 5 |
| GO:0098542 | defense response to other organism | BP | 0.004 | Cluster 5 |
| GO:0051707 | response to other organism | BP | 0.005 | Cluster 5 |
| GO:0002682 | regulation of immune system process | BP | 0.005 | Cluster 5 |
| GO:0061134 | peptidase regulator activity | MF | 0.007 | Cluster 5 |
| GO:0030451 | regulation of complement activation, alternative pathway | BP | 0.016 | Cluster 5 |
| GO:0030246 | carbohydrate binding | MF | 0.017 | Cluster 5 |
| GO:0002920 | regulation of humoral immune response | BP | 0.022 | Cluster 5 |
| GO:0042742 | defense response to bacterium | BP | 0.038 | Cluster 5 |
| GO:0043207 | response to external biotic stimulus | BP | 0.041 | Cluster 5 |
| GO:0018401 | peptidyl-proline hydroxylation to 4-hydroxy-L-proline | BP | <0.0001 | Cluster 6 |
| GO:0019471 | 4-hydroxyproline metabolic process | BP | <0.0001 | Cluster 6 |
| GO:0019511 | peptidyl-proline hydroxylation | BP | <0.0001 | Cluster 6 |
| GO:0018126 | protein hydroxylation | BP | <0.0001 | Cluster 6 |
| GO:0031545 | peptidyl-proline 4-dioxygenase activity | MF | <0.0001 | Cluster 6 |
| GO:0018208 | peptidyl-proline modification | BP | <0.0001 | Cluster 6 |
| GO:0031543 | peptidyl-proline dioxygenase activity | MF | <0.0001 | Cluster 6 |
| GO:0031418 | L-ascorbic acid binding | MF | <0.0001 | Cluster 6 |
| GO:0016706 | 2-oxoglutarate-dependent dioxygenase activity | MF | 0.0001 | Cluster 6 |
| GO:0005506 | iron ion binding | MF | 0.0005 | Cluster 6 |
| GO:0051213 | dioxygenase activity | MF | 0.0005 | Cluster 6 |
| GO:0006575 | cellular modified amino acid metabolic process | BP | 0.0005 | Cluster 6 |
| GO:0016705 | oxidoreductase activity, acting on paired donors, with incorporation or reduction of molecular oxygen | MF | 0.001 | Cluster 6 |
| GO:1901605 | alpha-amino acid metabolic process | BP | 0.001 | Cluster 6 |
| GO:0048029 | monosaccharide binding | MF | 0.003 | Cluster 6 |
| GO:0008198 | ferrous iron binding | MF | 0.005 | Cluster 6 |
| GO:0006520 | cellular amino acid metabolic process | BP | 0.005 | Cluster 6 |
| GO:0019842 | vitamin binding | MF | 0.006 | Cluster 6 |
| GO:0071456 | cellular response to hypoxia | BP | 0.008 | Cluster 6 |
| GO:0036294 | cellular response to decreased oxygen levels | BP | 0.011 | Cluster 6 |
| GO:0071453 | cellular response to oxygen levels | BP | 0.015 | Cluster 6 |
| GO:0016491 | oxidoreductase activity | MF | 0.028 | Cluster 6 |
| GO:0031406 | carboxylic acid binding | MF | 0.028 | Cluster 6 |

| | | | | |
|---|---|---|---|---|
| GO:0043177 | organic acid binding | MF | 0.030 | Cluster 6 |
| GO:0001937 | negative regulation of endothelial cell proliferation | BP | <0.0001 | Same dir. up |
| GO:0004051 | arachidonate 5-lipoxygenase activity | MF | <0.0001 | Same dir. up |
| GO:0031407 | oxylipin metabolic process | BP | <0.0001 | Same dir. up |
| GO:0031408 | oxylipin biosynthetic process | BP | <0.0001 | Same dir. up |
| GO:1901751 | leukotriene A4 metabolic process | BP | <0.0001 | Same dir. up |
| GO:1901753 | leukotriene A4 biosynthetic process | BP | <0.0001 | Same dir. up |
| GO:1904999 | positive regulation of leukocyte adhesion to arterial endothelial cell | BP | <0.0001 | Same dir. up |
| GO:0006959 | humoral immune response | BP | <0.0001 | Same dir. up |
| GO:0106014 | regulation of inflammatory response to wounding | BP | <0.0001 | Same dir. up |
| GO:0061044 | negative regulation of vascular wound healing | BP | <0.0001 | Same dir. up |
| GO:0097176 | epoxide metabolic process | BP | <0.0001 | Same dir. up |
| GO:0004052 | arachidonate 12(S)-lipoxygenase activity | MF | <0.0001 | Same dir. up |
| GO:0002232 | leukocyte chemotaxis involved in inflammatory response | BP | <0.0001 | Same dir. up |
| GO:0036403 | arachidonate 8(S)-lipoxygenase activity | MF | <0.0001 | Same dir. up |
| GO:2001301 | lipoxin biosynthetic process | BP | <0.0001 | Same dir. up |
| GO:1904996 | positive regulation of leukocyte adhesion to vascular endothelial cell | BP | <0.0001 | Same dir. up |
| GO:1904997 | regulation of leukocyte adhesion to arterial endothelial cell | BP | <0.0001 | Same dir. up |
| GO:2001300 | lipoxin metabolic process | BP | <0.0001 | Same dir. up |
| GO:0061043 | regulation of vascular wound healing | BP | <0.0001 | Same dir. up |
| GO:0002523 | leukocyte migration involved in inflammatory response | BP | <0.0001 | Same dir. up |
| GO:1904994 | regulation of leukocyte adhesion to vascular endothelial cell | BP | <0.0001 | Same dir. up |
| GO:1901503 | ether biosynthetic process | BP | <0.0001 | Same dir. up |
| GO:0005641 | nuclear envelope lumen | CC | <0.0001 | Same dir. up |
| GO:1903671 | negative regulation of sprouting angiogenesis | BP | <0.0001 | Same dir. up |
| GO:0018904 | ether metabolic process | BP | <0.0001 | Same dir. up |
| GO:0001676 | long-chain fatty acid metabolic process | BP | <0.0001 | Same dir. up |
| GO:0043651 | linoleic acid metabolic process | BP | <0.0001 | Same dir. up |
| GO:0042759 | long-chain fatty acid biosynthetic process | BP | <0.0001 | Same dir. up |
| GO:0016702 | oxidoreductase activity, acting on single donors with incorporation of molecular oxygen, incorporation of two atoms of | MF | <0.0001 | Same dir. up |

| | | | | |
|---|---|---|---|---|
| | oxygen | | | |
| GO:0019370 | leukotriene biosynthetic process | BP | <0.0001 | Same dir. up |
| GO:0016701 | oxidoreductase activity, acting on single donors with incorporation of molecular oxygen | MF | <0.0001 | Same dir. up |
| GO:0042383 | sarcolemma | CC | <0.0001 | Same dir. up |
| GO:0002540 | leukotriene production involved in inflammatory response | BP | <0.0001 | Same dir. up |
| GO:0002538 | arachidonic acid metabolite production involved in inflammatory response | BP | <0.0001 | Same dir. up |
| GO:0001936 | regulation of endothelial cell proliferation | BP | 0.0001 | Same dir. up |
| GO:0006691 | leukotriene metabolic process | BP | 0.0001 | Same dir. up |
| GO:0051121 | hepoxilin metabolic process | BP | 0.0002 | Same dir. up |
| GO:0051122 | hepoxilin biosynthetic process | BP | 0.0002 | Same dir. up |
| GO:1904734 | positive regulation of electron transfer activity | BP | 0.0002 | Same dir. up |
| GO:1904960 | positive regulation of cytochrome-c oxidase activity | BP | 0.0002 | Same dir. up |
| GO:0005506 | iron ion binding | MF | 0.0002 | Same dir. up |
| GO:0033559 | unsaturated fatty acid metabolic process | BP | 0.0002 | Same dir. up |
| GO:0006636 | unsaturated fatty acid biosynthetic process | BP | 0.0002 | Same dir. up |
| GO:0009605 | response to external stimulus | BP | 0.0002 | Same dir. up |
| GO:0061045 | negative regulation of wound healing | BP | 0.0002 | Same dir. up |
| GO:0002532 | production of molecular mediator involved in inflammatory response | BP | 0.0002 | Same dir. up |
| GO:0046456 | icosanoid biosynthetic process | BP | 0.0002 | Same dir. up |
| GO:0016525 | negative regulation of angiogenesis | BP | 0.0003 | Same dir. up |
| GO:0019369 | arachidonic acid metabolic process | BP | 0.0003 | Same dir. up |
| GO:0031970 | organelle envelope lumen | CC | 0.0003 | Same dir. up |
| GO:0005391 | P-type sodium:potassium-exchanging transporter activity | MF | 0.0003 | Same dir. up |
| GO:0008556 | P-type potassium transmembrane transporter activity | MF | 0.0003 | Same dir. up |
| GO:0019372 | lipoxygenase pathway | BP | 0.0003 | Same dir. up |
| GO:2000181 | negative regulation of blood vessel morphogenesis | BP | 0.0003 | Same dir. up |
| GO:0050680 | negative regulation of epithelial cell proliferation | BP | 0.0003 | Same dir. up |
| GO:1904732 | regulation of electron transfer activity | BP | 0.0003 | Same dir. up |
| GO:1904959 | regulation of cytochrome-c oxidase | BP | 0.0003 | Same dir. up |

| | | | | |
|---|---|---|---|---|
| | activity | | | |
| GO:0006690 | icosanoid metabolic process | BP | 0.0003 | Same dir. up |
| GO:1903035 | negative regulation of response to wounding | BP | 0.0004 | Same dir. up |
| GO:1903670 | regulation of sprouting angiogenesis | BP | 0.0004 | Same dir. up |
| GO:0042379 | chemokine receptor binding | MF | 0.0005 | Same dir. up |
| GO:1901343 | negative regulation of vasculature development | BP | 0.0005 | Same dir. up |
| GO:0019229 | regulation of vasoconstriction | BP | 0.0006 | Same dir. up |
| GO:0005576 | extracellular region | CC | 0.0007 | Same dir. up |
| GO:1903524 | positive regulation of blood circulation | BP | 0.0009 | Same dir. up |
| GO:1903573 | negative regulation of response to endoplasmic reticulum stress | BP | 0.001 | Same dir. up |
| GO:0008009 | chemokine activity | MF | 0.001 | Same dir. up |
| GO:0045907 | positive regulation of vasoconstriction | BP | 0.001 | Same dir. up |
| GO:0090084 | negative regulation of inclusion body assembly | BP | 0.001 | Same dir. up |
| GO:0030501 | positive regulation of bone mineralization | BP | 0.001 | Same dir. up |
| GO:0070169 | positive regulation of biomineral tissue development | BP | 0.002 | Same dir. up |
| GO:1901568 | fatty acid derivative metabolic process | BP | 0.002 | Same dir. up |
| GO:0002376 | immune system process | BP | 0.002 | Same dir. up |
| GO:1901570 | fatty acid derivative biosynthetic process | BP | 0.002 | Same dir. up |
| GO:0048878 | chemical homeostasis | BP | 0.002 | Same dir. up |
| GO:0006954 | inflammatory response | BP | 0.002 | Same dir. up |
| GO:0015079 | potassium ion transmembrane transporter activity | MF | 0.002 | Same dir. up |
| GO:0006955 | immune response | BP | 0.002 | Same dir. up |
| GO:0005215 | transporter activity | MF | 0.002 | Same dir. up |
| GO:0010155 | regulation of proton transport | BP | 0.002 | Same dir. up |
| GO:0061041 | regulation of wound healing | BP | 0.003 | Same dir. up |
| GO:0030595 | leukocyte chemotaxis | BP | 0.003 | Same dir. up |
| GO:1900407 | regulation of cellular response to oxidative stress | BP | 0.003 | Same dir. up |
| GO:0019233 | sensory perception of pain | BP | 0.003 | Same dir. up |
| GO:0045598 | regulation of fat cell differentiation | BP | 0.003 | Same dir. up |
| GO:0090083 | regulation of inclusion body assembly | BP | 0.004 | Same dir. up |
| GO:0015662 | P-type ion transporter activity | MF | 0.004 | Same dir. up |
| GO:0050678 | regulation of epithelial cell proliferation | BP | 0.004 | Same dir. up |
| GO:0042593 | glucose homeostasis | BP | 0.004 | Same dir. up |
| GO:0033500 | carbohydrate homeostasis | BP | 0.004 | Same dir. up |

| | | | | |
|---|---|---|---|---|
| GO:1903034 | regulation of response to wounding | BP | 0.005 | Same dir. up |
| GO:0030500 | regulation of bone mineralization | BP | 0.006 | Same dir. up |
| GO:0015081 | sodium ion transmembrane transporter activity | MF | 0.006 | Same dir. up |
| GO:0032412 | regulation of ion transmembrane transporter activity | BP | 0.006 | Same dir. up |
| GO:1902882 | regulation of response to oxidative stress | BP | 0.006 | Same dir. up |
| GO:0034440 | lipid oxidation | BP | 0.007 | Same dir. up |
| GO:0022898 | regulation of transmembrane transporter activity | BP | 0.007 | Same dir. up |
| GO:0006633 | fatty acid biosynthetic process | BP | 0.007 | Same dir. up |
| GO:0022857 | transmembrane transporter activity | MF | 0.007 | Same dir. up |
| GO:0006952 | defense response | BP | 0.008 | Same dir. up |
| GO:0055093 | response to hyperoxia | BP | 0.008 | Same dir. up |
| GO:0070167 | regulation of biomineral tissue development | BP | 0.008 | Same dir. up |
| GO:1905897 | regulation of response to endoplasmic reticulum stress | BP | 0.009 | Same dir. up |
| GO:0032409 | regulation of transporter activity | BP | 0.009 | Same dir. up |
| GO:0035296 | regulation of tube diameter | BP | 0.011 | Same dir. up |
| GO:0097746 | blood vessel diameter maintenance | BP | 0.011 | Same dir. up |
| GO:1903426 | regulation of reactive oxygen species biosynthetic process | BP | 0.011 | Same dir. up |
| GO:0035150 | regulation of tube size | BP | 0.011 | Same dir. up |
| GO:0050896 | response to stimulus | BP | 0.011 | Same dir. up |
| GO:0034765 | regulation of ion transmembrane transport | BP | 0.011 | Same dir. up |
| GO:0060351 | cartilage development involved in endochondral bone morphogenesis | BP | 0.011 | Same dir. up |
| GO:0031667 | response to nutrient levels | BP | 0.011 | Same dir. up |
| GO:0051353 | positive regulation of oxidoreductase activity | BP | 0.012 | Same dir. up |
| GO:2000377 | regulation of reactive oxygen species metabolic process | BP | 0.013 | Same dir. up |
| GO:0034762 | regulation of transmembrane transport | BP | 0.013 | Same dir. up |
| GO:0006631 | fatty acid metabolic process | BP | 0.013 | Same dir. up |
| GO:0032414 | positive regulation of ion transmembrane transporter activity | BP | 0.013 | Same dir. up |
| GO:0005615 | extracellular space | CC | 0.014 | Same dir. up |
| GO:0055078 | sodium ion homeostasis | BP | 0.014 | Same dir. up |
| GO:0045778 | positive regulation of ossification | BP | 0.014 | Same dir. up |
| GO:0036336 | dendritic cell migration | BP | 0.014 | Same dir. up |

| | | | | |
|---|---|---|---|---|
| GO:0009991 | response to extracellular stimulus | BP | 0.015 | Same dir. up |
| GO:0036296 | response to increased oxygen levels | BP | 0.015 | Same dir. up |
| GO:0001848 | complement binding | MF | 0.015 | Same dir. up |
| GO:0060326 | cell chemotaxis | BP | 0.015 | Same dir. up |
| GO:0001664 | G protein-coupled receptor binding | MF | 0.016 | Same dir. up |
| GO:0032411 | positive regulation of transporter activity | BP | 0.017 | Same dir. up |
| GO:1904062 | regulation of cation transmembrane transport | BP | 0.018 | Same dir. up |
| GO:0051213 | dioxygenase activity | MF | 0.018 | Same dir. up |
| GO:0006957 | complement activation, alternative pathway | BP | 0.018 | Same dir. up |
| GO:1900015 | regulation of cytokine production involved in inflammatory response | BP | 0.019 | Same dir. up |
| GO:0071805 | potassium ion transmembrane transport | BP | 0.019 | Same dir. up |
| GO:0001872 | (1->3)-beta-D-glucan binding | MF | 0.022 | Same dir. up |
| GO:0022853 | active ion transmembrane transporter activity | MF | 0.022 | Same dir. up |
| GO:1990573 | potassium ion import across plasma membrane | BP | 0.023 | Same dir. up |
| GO:0002526 | acute inflammatory response | BP | 0.024 | Same dir. up |
| GO:0072330 | monocarboxylic acid biosynthetic process | BP | 0.024 | Same dir. up |
| GO:1903522 | regulation of blood circulation | BP | 0.025 | Same dir. up |
| GO:0019829 | ATPase-coupled cation transmembrane transporter activity | MF | 0.028 | Same dir. up |
| GO:0043269 | regulation of ion transport | BP | 0.028 | Same dir. up |
| GO:0008285 | negative regulation of cell population proliferation | BP | 0.028 | Same dir. up |
| GO:0006812 | cation transport | BP | 0.028 | Same dir. up |
| GO:0050900 | leukocyte migration | BP | 0.029 | Same dir. up |
| GO:0030299 | intestinal cholesterol absorption | BP | 0.029 | Same dir. up |
| GO:1904064 | positive regulation of cation transmembrane transport | BP | 0.030 | Same dir. up |
| GO:0003013 | circulatory system process | BP | 0.031 | Same dir. up |
| GO:0003018 | vascular process in circulatory system | BP | 0.031 | Same dir. up |
| GO:0051341 | regulation of oxidoreductase activity | BP | 0.031 | Same dir. up |
| GO:0009725 | response to hormone | BP | 0.032 | Same dir. up |
| GO:0005125 | cytokine activity | MF | 0.032 | Same dir. up |
| GO:0055075 | potassium ion homeostasis | BP | 0.033 | Same dir. up |
| GO:0006813 | potassium ion transport | BP | 0.035 | Same dir. up |
| GO:0042625 | ATPase-coupled ion transmembrane transporter activity | MF | 0.041 | Same dir. up |
| GO:0034767 | positive regulation of ion transmembrane | BP | 0.042 | Same dir. up |

| | | | | |
|---|---|---|---|---|
| | transport | | | |
| GO:0042592 | homeostatic process | BP | 0.043 | Same dir. up |
| GO:0043005 | neuron projection | CC | 0.044 | Same dir. up |
| GO:0030007 | cellular potassium ion homeostasis | BP | 0.044 | Same dir. up |
| GO:0005890 | sodium:potassium-exchanging ATPase complex | CC | 0.044 | Same dir. up |
| GO:0070852 | cell body fiber | CC | 0.044 | Same dir. up |
| GO:0016363 | nuclear matrix | CC | 0.044 | Same dir. up |
| GO:0034764 | positive regulation of transmembrane transport | BP | 0.044 | Same dir. up |
| GO:0098856 | intestinal lipid absorption | BP | 0.044 | Same dir. up |
| GO:0051087 | chaperone binding | MF | 0.048 | Same dir. up |
| GO:0046873 | metal ion transmembrane transporter activity | MF | 0.048 | Same dir. up |

Soft clustering was performed to visualize differential expression patterns and clusters that showed expression patterns with differential expression between H-DO, Imm. and L-DO, Imm. samples that is no longer differentially expressed in the L-DO, Acc. samples were run in the gene ontology analysis (Cluster 5 and 6). Differentially expressed genes identified in the L-DO, Imm. vs L-DO, Acc. comparison that were identified as being expressed in the same direction in the L-DO, Acc. and H-DO, Imm. samples (Same dir. up) were also put through gene ontology analysis.

**Supplemental Table 6. Significant GO terms from gene ontology analysis on *E. neumayeri*.**

| GO-term | Name | Ontology | *q*-value | Expression Pattern |
|---|---|---|---|---|
| GO:0032364 | oxygen homeostasis | BP | 0.002 | Cluster 5 |
| GO:0033483 | gas homeostasis | BP | 0.002 | Cluster 5 |
| GO:0051344 | negative regulation of cyclic-nucleotide phosphodiesterase activity | BP | 0.002 | Cluster 5 |
| GO:0099159 | regulation of modification of postsynaptic structure | BP | 0.002 | Cluster 5 |
| GO:0005201 | extracellular matrix structural constituent | MF | 0.002 | Cluster 5 |
| GO:0140252 | regulation protein catabolic process at postsynapse | BP | 0.003 | Cluster 5 |
| GO:0002412 | antigen transcytosis by M cells in mucosal-associated lymphoid tissue | BP | 0.003 | Cluster5 |
| GO:0051342 | regulation of cyclic-nucleotide phosphodiesterase activity | BP | 0.005 | Cluster 5 |
| GO:0060711 | labyrinthine layer development | BP | 0.006 | Cluster 5 |
| GO:0031545 | peptidyl-proline 4-dioxygenase activity | MF | 0.007 | Cluster 5 |
| GO:0060347 | heart trabecula formation | BP | 0.008 | Cluster 5 |
| GO:0031543 | peptidyl-proline dioxygenase activity | MF | 0.008 | Cluster 5 |

| | | | | |
|---|---|---|---|---|
| GO:0018401 | peptidyl-proline hydroxylation to 4-hydroxy-L-proline | BP | 0.009 | Cluster 5 |
| GO:0008198 | ferrous iron binding | MF | 0.012 | Cluster 5 |
| GO:0019471 | 4-hydroxyproline metabolic process | BP | 0.012 | Cluster 5 |
| GO:0019511 | peptidyl-proline hydroxylation | BP | 0.012 | Cluster 5 |
| GO:0001666 | response to hypoxia | BP | 0.012 | Cluster 5 |
| GO:1905290 | negative regulation of CAMKK-AMPK signaling cascade | BP | 0.012 | Cluster 5 |
| GO:0071731 | response to nitric oxide | BP | 0.012 | Cluster 5 |
| GO:0060343 | trabecula formation | BP | 0.012 | Cluster 5 |
| GO:0031418 | L-ascorbic acid binding | MF | 0.012 | Cluster 5 |
| GO:0055008 | cardiac muscle tissue morphogenesis | BP | 0.012 | Cluster 5 |
| GO:0036293 | response to decreased oxygen levels | BP | 0.012 | Cluster 5 |
| GO:0045056 | transcytosis | BP | 0.015 | Cluster 5 |
| GO:0060415 | muscle tissue morphogenesis | BP | 0.015 | Cluster 5 |
| GO:1905289 | regulation of CAMKK-AMPK signaling cascade | BP | 0.016 | Cluster 5 |
| GO:0042589 | zymogen granule membrane | CC | 0.016 | Cluster 5 |
| GO:0070482 | response to oxygen levels | BP | 0.016 | Cluster 5 |
| GO:0018126 | protein hydroxylation | BP | 0.020 | Cluster 5 |
| GO:0005344 | oxygen carrier activity | MF | 0.026 | Cluster 5 |
| GO:0071456 | cellular response to hypoxia | BP | 0.033 | Cluster 5 |
| GO:0060412 | ventricular septum morphogenesis | BP | 0.044 | Cluster 5 |
| GO:0036294 | cellular response to decreased oxygen levels | BP | 0.046 | Cluster 5 |
| GO:0019825 | oxygen binding | MF | 0.046 | Cluster 5 |
| GO:0018208 | peptidyl-proline modification | BP | 0.046 | Cluster 5 |
| GO:0005506 | iron ion binding | MF | 0.049 | Cluster 5 |

Soft clustering was performed to visualize differential expression patterns and clusters that showed expression patterns with differential expression between H-DO, Imm. and L-DO, Imm. samples that are no longer differentially expressed in the L-DO, Acc. samples were run in the gene ontology analysis (Cluster 5). Differentially expressed genes identified in the L-DO, Imm. vs L-DO, Acc. comparison that were identified as being expressed in the same direction in the L-DO, Acc. and H-DO, Imm. samples were also put through gene ontology analysis, however, there were no significant results.

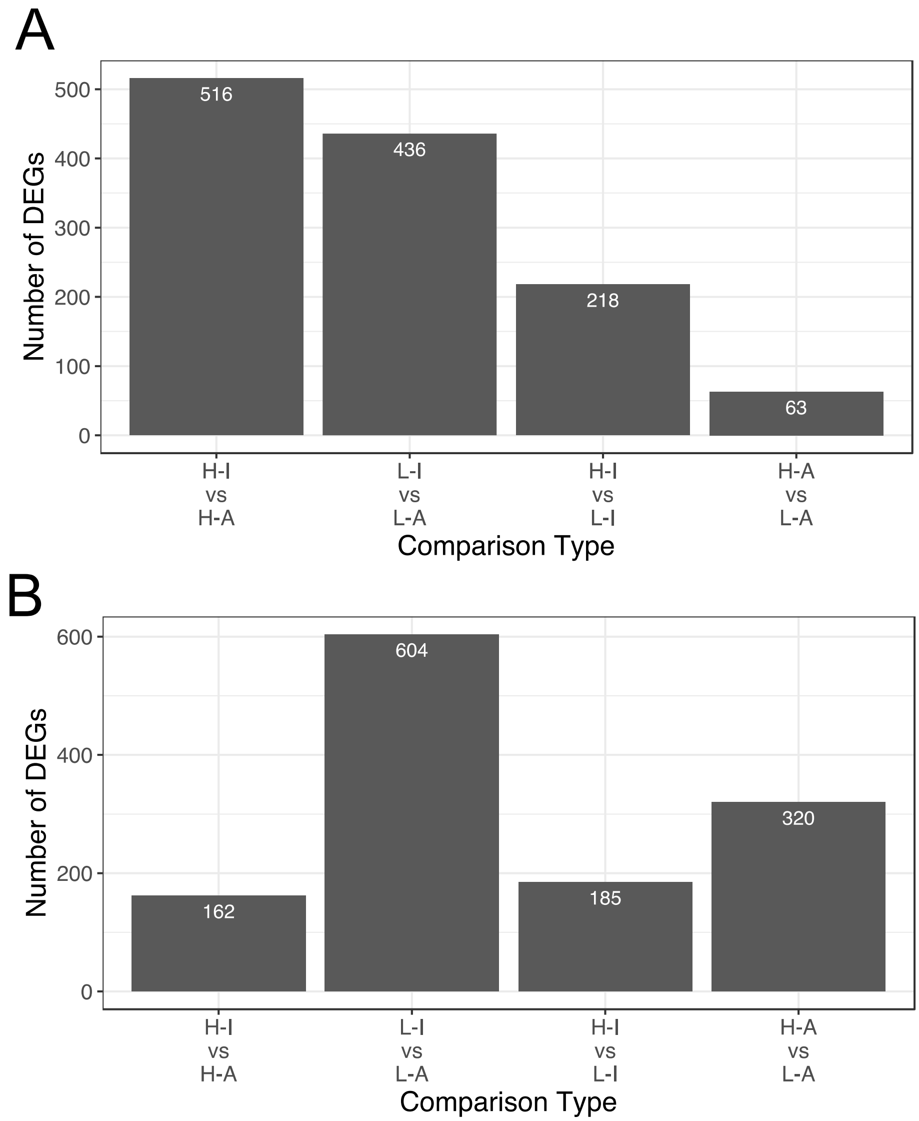


**Supplemental Figure 1. Number of differentially expressed genes identified for each type of comparison made for (A) *E. neumayeri* and (B) *E. apleurogramma*.** L-DO = low DO source, H-DO = high DO source, I. = immediately sampled, A. = sampled after acclimation.

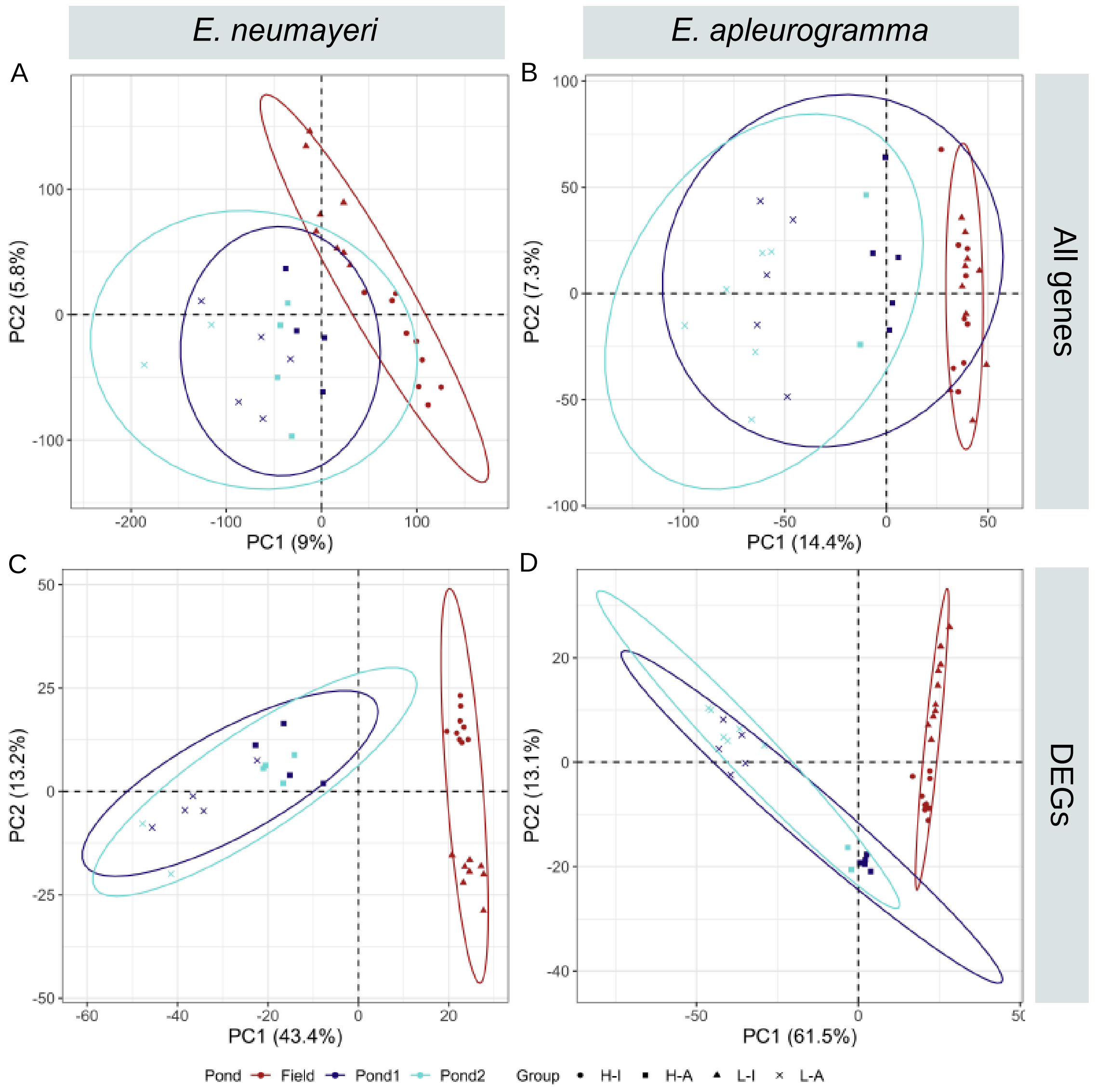


**Supplemental Figure 2. Principle component analysis (PCA) on TMM normalized, log2 transformed, and median centered gene expression values for (A and B) all genes and (C and D) all differentially expressed genes in *E. neumayeri* (A and C) and *E. apleurogramma* (B and D) labelled with acclimation pond.** PCA shows clustering by sample type but no clustering by acclimation pond. L-DO = low DO source, H-DO = high DO source, I. = immediately sampled, A. = sampled after acclimation.

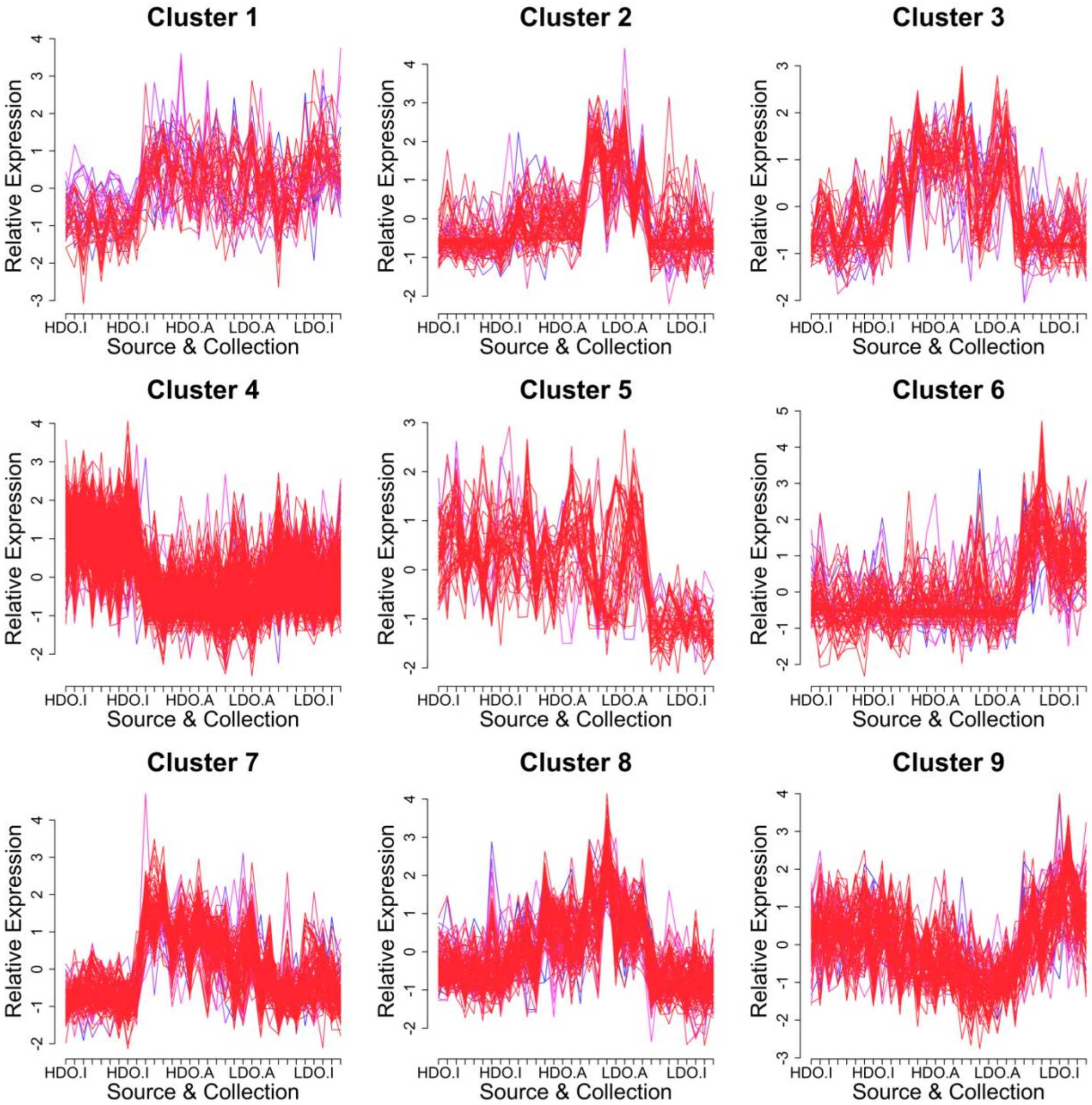


**Supplemental Figure 3. Soft cluster results for *E. neumayeri*.** Soft clustering allows for the calculation of a membership score for every gene in each cluster. Only genes with a membership score > 0.7 were plotted. Red lines indicate genes with a membership score > 0.9, purple lines have a membership score < 0.9 and > 0.8, and blue lines have a membership score < 0.8 and > 0.7. L-DO = low DO source, H-DO = high DO source, I. = immediately sampled, A. = sampled after acclimation. Clusters 5 and 6 were selected for gene ontology (GO) analysis.

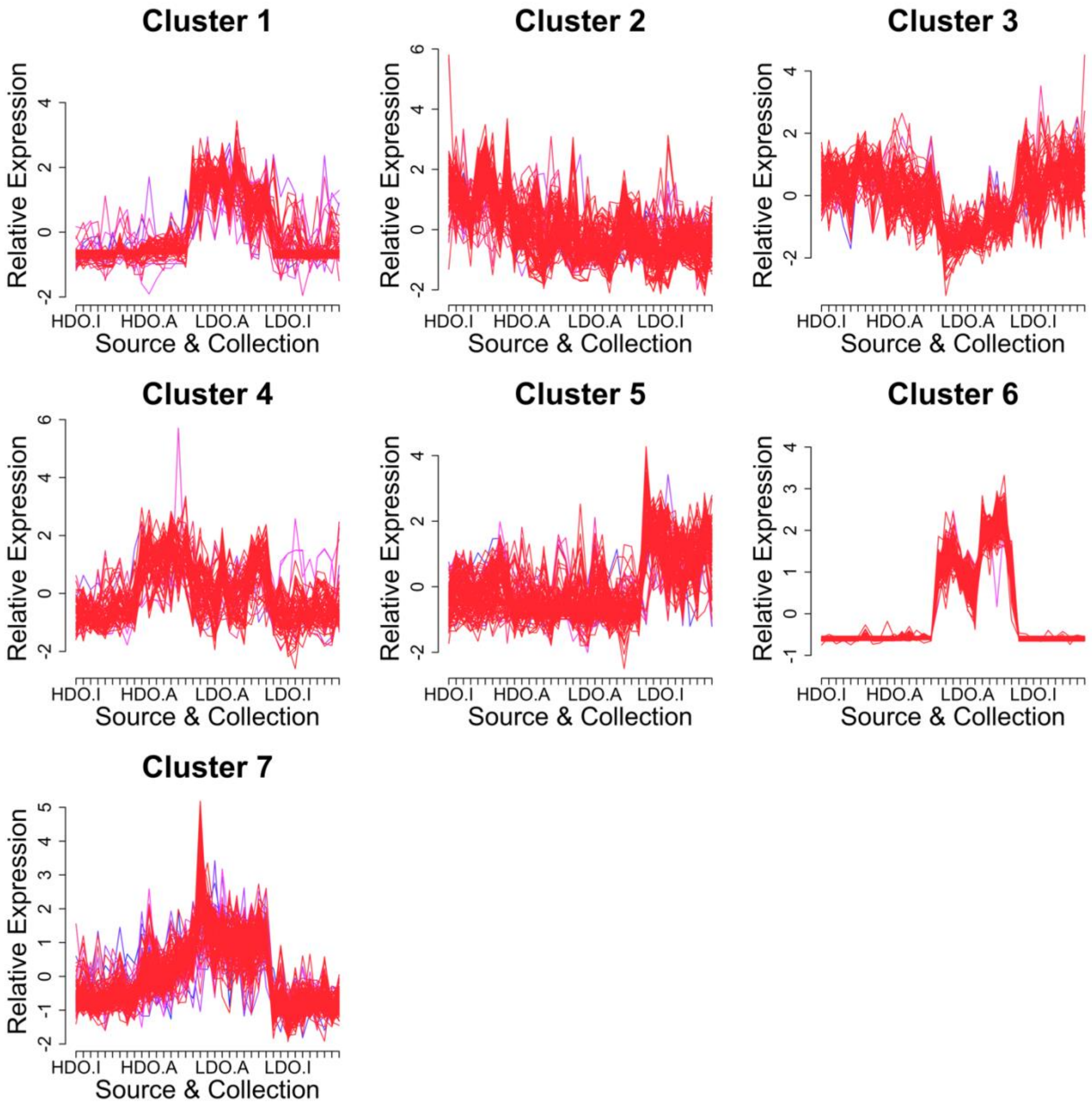


**Supplemental Figure 4. Soft cluster results for *E. apleurogramma*.** Soft clustering allows for the calculation of a membership score for every gene in each cluster. Only genes with a membership score > 0.7 were plotted. Red lines indicate genes with a membership score > 0.9, purple lines have a membership score < 0.9 and > 0.8, and blue lines have a membership score < 0.8 and > 0.7. L-DO = low DO source, H-DO = high DO source, I. = immediately sampled, A. = sampled after acclimation. Cluster 5 was selected for gene ontology (GO) analysis.

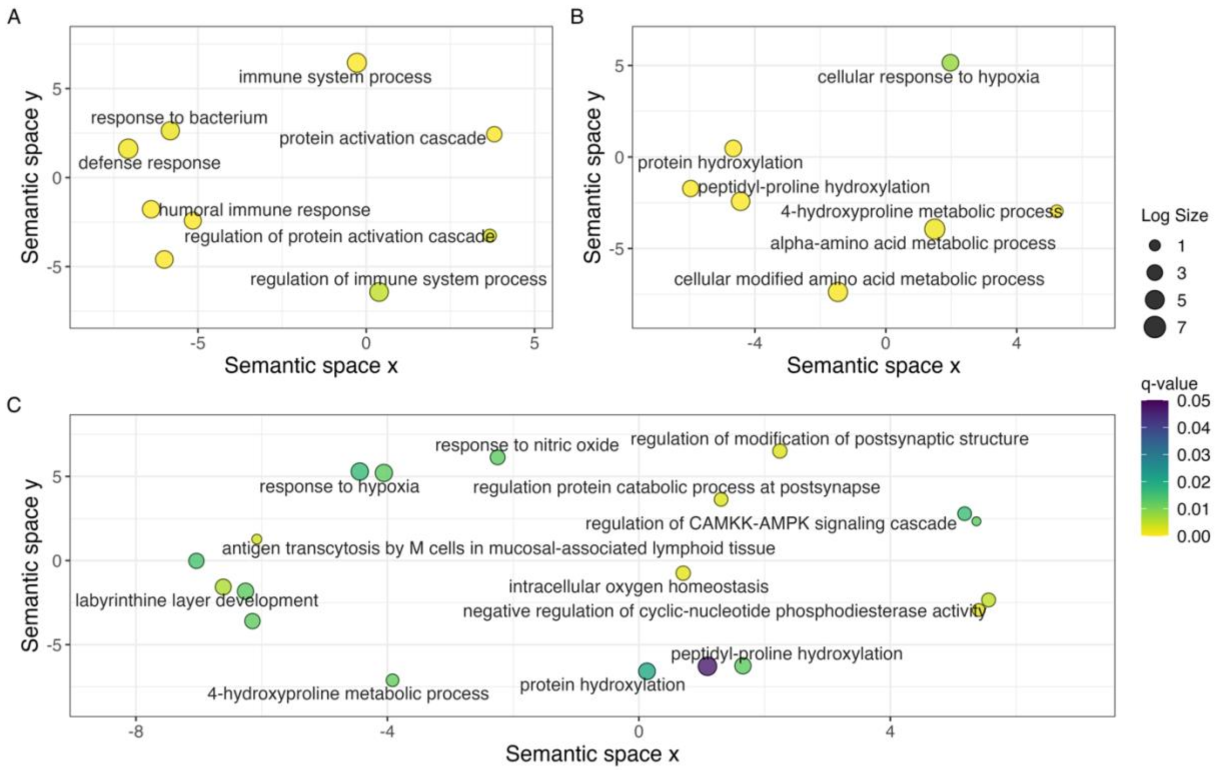


**Supplemental Figure 5. REVIGO analysis of gene ontology results for *E. neumayeri* (native) and *E. apleurogramma* (range-expanding).** Gene ontology enrichment analysis was run on clusters of interest identified from the soft clustering analysis. Results were then analyzed using REVIGO which removes redundant GO terms and performs SimRel clustering to plot the similarity of given GO terms in semantic space. Circle size indicates the number of GO child terms and the color of the circle shows the q value with yellow representing $q < 0.001$. (A) Cluster 5 from *E. neumayeri* represents genes with decreased expression in L-I samples compared to the rest of the samples. (B) Cluster 6 from *E. neumayeri* and (C) cluster 5 from *E. apleurogramma* contains genes with increased expression in L-I samples compared to the rest of the samples.
